# Plant MutS2 proteins function in plastid ribosome quality control

**DOI:** 10.1101/2025.09.02.673837

**Authors:** Amanda K. Broz, Kalia Kodrich, Kasavajhala V.S.K. Prasad, Shady A. Kuster, Sunkyung Lee, Evan S. Forsythe, Daniel B. Sloan

## Abstract

Tight regulation of chloroplast translation is essential for plant growth, development and environmental response. Active translation can result in stalling and collision of ribosomes, which have negative fitness consequences. However, the ways in which chloroplasts respond to these types of translational stressors remain unknown. Here, we identify two MutS2 proteins that act as critical players in plastid ribosome-associated quality control (RQC) in Arabidopsis. We found that both MutS2A and MutS2B are required to overcome specific antibiotic-induced ribosome stalling and collisions. Further, these proteins appear to be essential for tissue greening during de-etiolation, potentially due to increased translational demand during the transition from etioplast to chloroplast. Although bacterial homologs of MutS2 have been widely recognized for their role in regulating homologous recombination, we found only weak support for this function in Arabidopsis plastids. Therefore, these proteins, which are widely conserved among photosynthetic eukaryotes, appear to be central in the resolution of ribosome collisions and may play a critical role during times of increased translational demand.

## Introduction

Chloroplasts (photosynthetic plastids) are a defining feature of plant cells and house the biochemical machinery that converts light energy into chemical energy. Plastids were acquired through endosymbiosis of a formerly free-living cyanobacteria and still retain their own genomes, which code for components of photosynthetic enzyme complexes as well as plastid transcriptional and translational machinery. Many nuclear-encoded genes are also critical for plastid function, and thus, tight coordination of these two genomes is required for plant growth, development, and response to environmental conditions. Even though plastid genomes contain fewer than 100 protein-coding genes, they account for the majority of mRNA transcripts in plant leaf tissue (∼75% in Arabidopsis) (Forsythe et al. 2022). Mature chloroplasts are found at high numbers (∼20-90) in plant leaf cells (Greiner et al. 2020), and chloroplast proteins account for a staggering ∼80% of the total protein in mesophyll cells (Heinemann et al. 2021). Thus, much of the transcriptional and translational burden of the photosynthetic process falls upon machinery present in the chloroplast.

The bulk of the work on chloroplast proteostasis has focused on plastid transcription (Puthiyaveetil et al. 2021; Zhang et al. 2023) and proteolysis (Sakamoto 2006; Nishimura et al. 2017; van Wijk 2024), with less attention towards plastid translation (Zoschke and Bock 2018). However, important advances in understanding chloroplast translation have been made, including the elucidation of the structure of plastid ribosomes (Bieri et al. 2017; Graf et al. 2017) and the identification of chloroplast RNA binding proteins that impact RNA stability and translation rates (Barkan and Small 2014). Ribosome pausing is prevalent along chloroplast mRNAs, and depends on mRNA sequence and secondary structure, as well as the sequence of the nascent peptide (Gawroński et al. 2018). Although these pauses are hypothesized to aid in the synthesis and assembly of the multi-subunit photosystems (Gawroński et al. 2018), a prolonged pause (ribosome stalling) is detrimental and can result in ribosome collisions (Collart and Weiss 2020).

In both prokaryotic and eukaryotic translation, ribosome stalling is increased under stress conditions and can be triggered by the presence of mRNAs that lack a stop codon (non-stop mRNAs), chemically damaged nucleotides, a lack of sufficient aminoacyl-tRNAs, or (in bacteria) various antibiotics (Moore and Sauer 2007; Wilson 2014; Yan and Zaher 2021; Nanjaraj Urs et al. 2024). When an mRNA is being translated by multiple ribosomes, stalling of the leading ribosome can result in collisions as the lagging ribosome(s) ‘catch up’. Collisions can have detrimental effects, as they tie up ribosomes, and the partial polypeptides created can be toxic to cells (Moore and Sauer 2007). Ribosome stalling can be prevalent during translation even under benign conditions. It has been previously reported that mRNAs are incorrectly processed at a rate of 0.4% in *Escherichia coli* (Moore and Sauer 2005), and between 6-20% of ribosomes are stalled as disomes (two collided ribosomes) in *Saccharomyces cerevisiae* (Diament et al. 2018; Zhao et al. 2021).

Parallel pathways have evolved to remedy ribosome stalling and collisions (Lytvynenko et al. 2019; Filbeck et al. 2022). In bacteria, the most well-characterized system is “trans-translation,” which uses a transfer-messenger RNA (tmRNA) to rescue ribosomes stalled on non-stop mRNAs (Moore and Sauer 2007). Here, the tmRNA inserts into the empty A site of the ribosome and is translated, adding a degron sequence to the nascent polypeptide after which ribosomes are recycled to prepare for another round of translation. In the absence of trans-translation, alternative ribosome rescue factors (ArfA and/or ArfB) can function in an additional ribosome rescue pathway. Arf enzymes share a structure similar to ribosome release factors (RFs) and bind to the empty A site on ribosomes paused on non-stop mRNAs, leading to hydrolysis of the peptidyl-tRNA. In this pathway, no labeling of the nascent polypeptide occurs, so it does not appear to play a role in protein quality control (Nadler et al. 2022).

In the cytosol of eukaryotes, stalled/collided ribosomes are initially sensed by “rescue factors” (Pelota/Dom and ABCE1/Rli1) that split the leading ribosome into its two subunits, initiating the ribosome-associated quality control (RQC) pathway (Filbeck et al. 2022). In RQC, the large ribosomal subunit (60S) retains the aberrant nascent polypeptide and is recognized by Rqc2/NEMF, which recruits the E3 ligase Listerin, promoting ubiquitination and subsequent degradation of the polypeptide (Bengtson and Joazeiro 2010). Homologs of Rqc2 (named RqcH) occur in ∼30% of bacterial genomes (Callan et al. 2024), but only recently was it discovered that RqcH and other proteins participate in a bacterial RQC pathway (Lytvynenko et al. 2019; Cerullo et al. 2022; Park et al. 2024).

In *Bacillus subtilis*, MutS2 was identified as a rescue factor in ribosome collisions, binding to and cleaving apart the subunits of the leading ribosome, thus initiating downstream RQC processes (Cerullo et al. 2022; Filbeck et al. 2022; Park et al. 2024). This novel role was surprising, as MutS2 is a member of the broader MutS family of proteins which are widely known for their roles in DNA recombination and repair (Malik and Henikoff 2000; Ogata et al. 2011). In fact, MutS2 was previously found to promote homologous recombination (HR) in *B. subtilis* (Burby and Simmons 2017). However, MutS2 suppresses HR in *Thermus thermophilus* (Fukui et al. 2008), *Heliobacter pylori* (Pinto et al. 2005; Damke et al. 2015), and *Thermotoga maritima* (Jeong et al. 2012).

MutS2, like other members of the MutS family, contains the core MutS domains (III-IV), followed by the highly conserved C-terminal domain V, which acts as an ATPase and is important in homodimerization (Fukui et al. 2004; Ogata et al. 2011). However, unlike the well characterized MutS1 from *E. coli,* MutS2 lacks an N-terminal mismatch recognition domain, and there is no evidence that it functions in mismatch repair (Eisen 1998; Malik and Henikoff 2000). In addition, MutS2 contains a unique C-terminal extension that encodes a Small MutS-related (Smr) domain that acts as a nicking endonuclease in *T. thermophilus* and *T. maritima* (Fukui et al. 2004, 2007; Jeong et al. 2012) and is proposed to cleave and resolve early recombination intermediates (Fukui et al. 2008; Fukui and Kuramitsu 2011).

Although the role of MutS2 in resolving ribosome collisions has only been extensively studied in *B. subtilis* (Cerullo et al. 2022; Park et al. 2024), this protein is found in diverse bacterial taxa (Callan et al. 2024; Park et al. 2024). Recent structural evidence from *T. thermophilus* suggests that cleavage of the leading ribosomal subunits is driven by an ATP-dependent conformational change in MutS2 (Fukui et al. 2025). In *B. subtilis*, the Smr domain of MutS2 was found to be essential for recruitment to stalled ribosomes (Park et al. 2024) and was originally hypothesized to cleave associated mRNAs (Cerullo et al. 2022). However, there is currently no evidence that it cleaves mRNAs associated with ribosome collisions (Park et al. 2024) even though the Smr domain has a structure similar to a ribonuclease (Fukui and Kuramitsu 2011). This was unexpected as other Smr domain-containing proteins (i.e., yeast CUE2 and *E. coli* SmrB) have been shown to cleave mRNAs associated with stalled ribosomes, facilitating rescue through the canonical pathways (D’Orazio et al. 2019; Saito et al. 2022).

Although *MutS2* is commonly found in bacterial genomes, it also occurs as a nuclear-encoded gene in photosynthetic eukaryotes (Fukui et al. 2004; Lin et al. 2007; Ogata et al. 2011; Berdieva et al. 2024; Sloan et al. 2024). Initial phylogenetic analyses revealed that plant *MutS2* is most closely related to *MutS2* from cyanobacteria, the plastid progenitor, suggesting an endosymbiotic gene transfer from the plastid to the nucleus (Lin et al. 2007). More recent studies in algae identified a second type of eukaryotic MutS2 protein that is more closely related to deltaproteobacteria and found in “CASH” (cryptophytes, alveolates, stramenopiles, haptophytes, and chlorarachniophytes) taxa (Berdieva et al. 2024).

Two widely conserved copies of *MutS2* are present within land plants and green algae, suggesting a gene duplication event that preceded the divergence of Viridiplantae (Lin et al. 2007; Sloan et al. 2024). In Arabidopsis and maize, both MutS2A and MutS2B have been found in plastid proteome fractions (Olinares et al. 2010; Majeran et al. 2012; Huang et al. 2013) and fusing Arabidopsis MutS2 transit peptides to GFP or RFP results in localization to the plastid (Carrie et al. 2009). Both genes are highly expressed across green tissues based on both mRNA (Waese et al. 2017) and protein abundance (van Wijk et al. 2021). However, in contrast to the extensive investigation of MutS2 in bacteria, the function of these genes in plants or any other eukaryote has remained entirely uncharacterized. Here, we examine the role of Arabidopsis MutS2 proteins in plastid RQC as well as HR in plastid genomes. We find only weak support for the involvement of MutS2 in plastid genome recombination. However, we provide evidence that both MutS2A and MutS2B are required to overcome antibiotic-induced ribosome stalling, suggesting that they play non-redundant roles in the plastid RQC pathway. We also show that Arabidopsis MutS2 proteins are essential for de-etiolation response (tissue greening and chloroplast differentiation following dark treatment), indicating that MutS2 RQC functions may be especially important during periods of intense translational demands in plastids.

## Results

### The origins, targeting, and molecular evolution of MutS2 in plants and algae

Phylogenetic analysis of MutS2 proteins from numerous taxa supports previous results that plant MutS2 proteins are of cyanobacterial origin and underwent an ancient duplication (Fig. 1, Supp. Fig. 1) (Lin et al. 2007; Sloan et al. 2024). One MutS2 homolog (MutS2B) retains the domain composition of bacterial MutS2, while the other (MutS2A) lacks the Smr endonuclease domain. All major green lineages appear to have retained both copies, whereas the sampled red algae and glaucophytes only have a single copy (Fig. 1).

**Figure 1.**
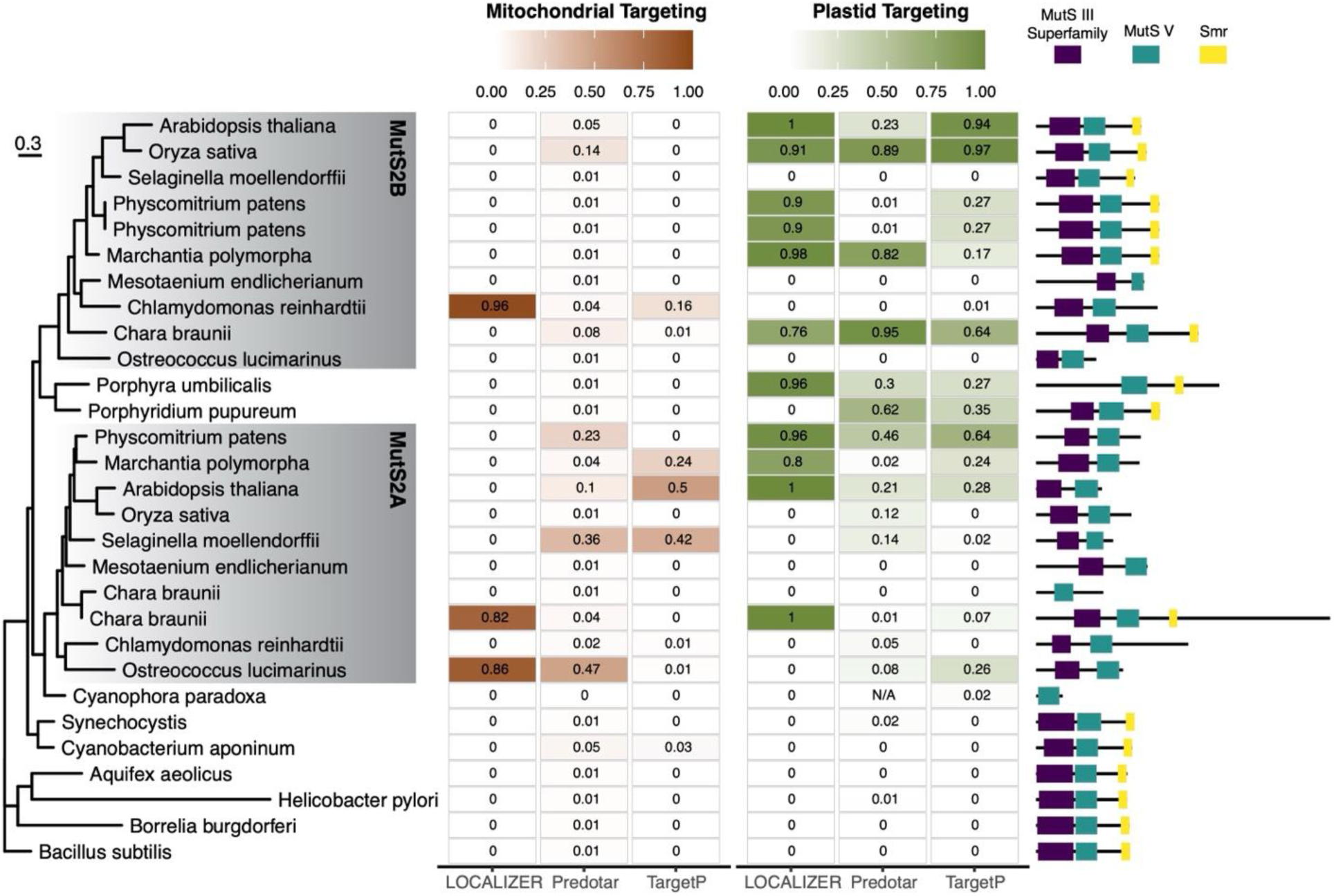
Phylogenetic, targeting, and structural analysis of MutS2 proteins. Green algae and land plants share two anciently duplicated copies of cyanobacterial-like MutS2 (gray boxes in the maximum-likelihood tree in the left panel). Bootstrap support values and sequence accessions for this tree are available in Supp. Fig. S1. The middle panel shows probabilities of mitochondrial and plastid targeting as predicted by three different in silico algorithms. The right panel shows the predicted domain architecture of each protein.

In silico targeting analyses of the MutS2 sequences from this diverse phylogenetic sample generally show stronger signals for localization in the plastid than in mitochondria (Fig. 1), which aligns with current evidence that both MutS2A and MutS2B are plastid-localized in angiosperms (Carrie et al. 2009; Olinares et al. 2010; Majeran et al. 2012; Huang et al. 2013). However, the predictions were not consistent across species, and many proteins did not show strong predictions for targeting to either organelle.

Domain architecture was relatively well conserved within the MutS2A and MutS2B groups, with the Smr domain typically retained in MutS2B but not in MutS2A. However, there are some exceptions.

For example, a MutS2A-like protein in the streptophytic green alga *Chara braunii* contains a predicted Smr domain despite the absence of that domain in the rest of the MutS2A clade. There were also proteins missing one or more of their expected domains (Fig. 1), including the absence of the MutS2B Smr domain in the sampled chlorophytes (Berdieva et al. 2024). However, any predictions of domain absence should be interpreted with caution, as they are sensitive to the accuracy of annotated gene models in the publicly available genomes used in the analysis.

The mature Arabidopsis MutS2A and MutS2B proteins share only ∼36% amino acid identity (after removal of gaps and the predicted transit peptide sequences) – approximately the same percent identity they share with their bacterial counterparts. The two have very different gene structures.

MutS2A (At5g54090.1) is comprised of 18 exons, whereas MutS2B (At1g65070.2) contains five exons. MutS2B has four reported splice isoforms, only one of which (At1g65070.2) encodes a full-length Smr domain. This isoform is the most highly expressed based on RNA-seq and expressed sequence tag (EST) data from The Arabidopsis Information Resource (Rhee et al. 2003)

### MutS2A and MutS2B show evidence of coexpression but not evolutionary rate covariation

Coexpression analysis can identify genes that share functional relationships or interactions. Using the pairwise coexpression viewer in the ATTED-II database (Obayashi et al. 2022), we found that MutS2A and MutS2B have correlated patterns of transcript abundance across tissue types and experimental conditions. Both genes fall within each other’s top 1% of most highly coexpressed genes. We subjected the top 300 coexpressed genes for both MutS2A (Supp. Table S1) and MutS2B (Supp. Table S2) to Gene Ontology (GO) enrichment analysis using PANTHER (version 19) (Mi et al. 2019; Thomas et al. 2022) (Supp. Table S3). In our analysis, 49 of the enriched GO terms were present in both MutS2A and MutS2B, while 33 were unique to MutS2A and 55 were unique to MutS2B (for a total of 137 enriched GO terms). For both MutS2A and MutS2B, cellular component and biological process categories showed enrichment for GO terms associated with the chloroplast (7 GO terms). In addition, enrichment in the biological process and molecular function categories included numerous GO terms involved in RNA binding and metabolism (44), DNA metabolism (17), nucleic acid metabolism (13) and translation (10). Biological processes involved in plant development were also enriched (11 GO terms).

Evolutionary rate covariation (ERC) is a technique that can identify proteins that physically interact or function in the same biochemical pathway based on the principle that co-functional genes face similar selection pressures, resulting in correlated increases and decreases in evolutionary rates across a phylogeny (Clark et al. 2012; Forsythe et al. 2025). Using our previously compiled ERC dataset from 20 angiosperm species (Forsythe et al. 2021), we found no signature of ERC between MutS2A and MutS2B (Pearson correlation; p = 0.70; R^2^ = 0.015). Analysis of MutS2A showed no significant ERC hits throughout the genome, while MutS2B returned seven significant hits (Supp. Table S4). This set did not show any obvious patterns related to function or subcellular localization, with one protein targeted to mitochondria, one to the peroxisome, one to the nucleus, and the remaining four to the cytosol. The overall lack of clear functional signature from MutS2B ERC hits makes it difficult to distinguish true co-functional evolution from noise.

### mutS2 mutants are sensitive to antibiotics that stall plastid ribosomes in an unrotated state

We took a genetic approach to understand the function of MutS2 in Arabidopsis and generated a segregating population of wild type (wt) individuals and single (*mutS2A* and *mutS2B*) and double (*mutS2A/B*) homozygous T-DNA mutants for comparative analyses. Lack of full-length gene expression in mutant lines was confirmed by reverse transcriptase PCR (Supp. Fig. S2). Under normal growth conditions, single and double *mutS2* mutants showed no apparent phenotype, paralleling results found in bacterial species that have shown MutS2 is not required for growth under normal laboratory conditions (Burby and Simmons 2017; Cerullo et al. 2022; Park et al. 2024).

To test for a role of MutS2 proteins in plastid RQC, we next determined whether these lines were sensitive to antibiotics that target plastid ribosomes. Due to the endosymbiotic origins of plastids, their ribosomes share many similarities with bacteria (Bieri et al. 2017) and are therefore typically inhibited by antibiotics that target bacterial translation (Oelmüller et al. 1986; Mulo et al. 2003). The macrolide antibiotic erythromycin inhibits nascent chain elongation by binding the exit site of the 50S ribosomal subunit in bacteria (Bulkley et al. 2010; Wilson 2014) and inhibits growth of *B. subtilis mutS2* mutants (Cerullo et al. 2022; Park et al. 2024). The impacts of erythromycin have not been studied extensively in Arabidopsis. However, studies in *Pisum sativum* and *Brassica campestris* have shown that erythromycin treatment inhibits photosynthetic efficiency and leads to reduced expression of plastid-encoded proteins, presumably through binding the plastid ribosome (Mulo et al. 2003; Yoon et al. 2020). By growing wt and *mutS2* mutants on media containing varying concentrations of erythromycin, we found that even low concentrations (0.4 μg/mL) had significant negative impacts on photosynthetic efficiency (Fv/Fm) and root growth in both single and double *mutS2* mutants when compared to wt controls (Fig. 2).

**Figure 2.**
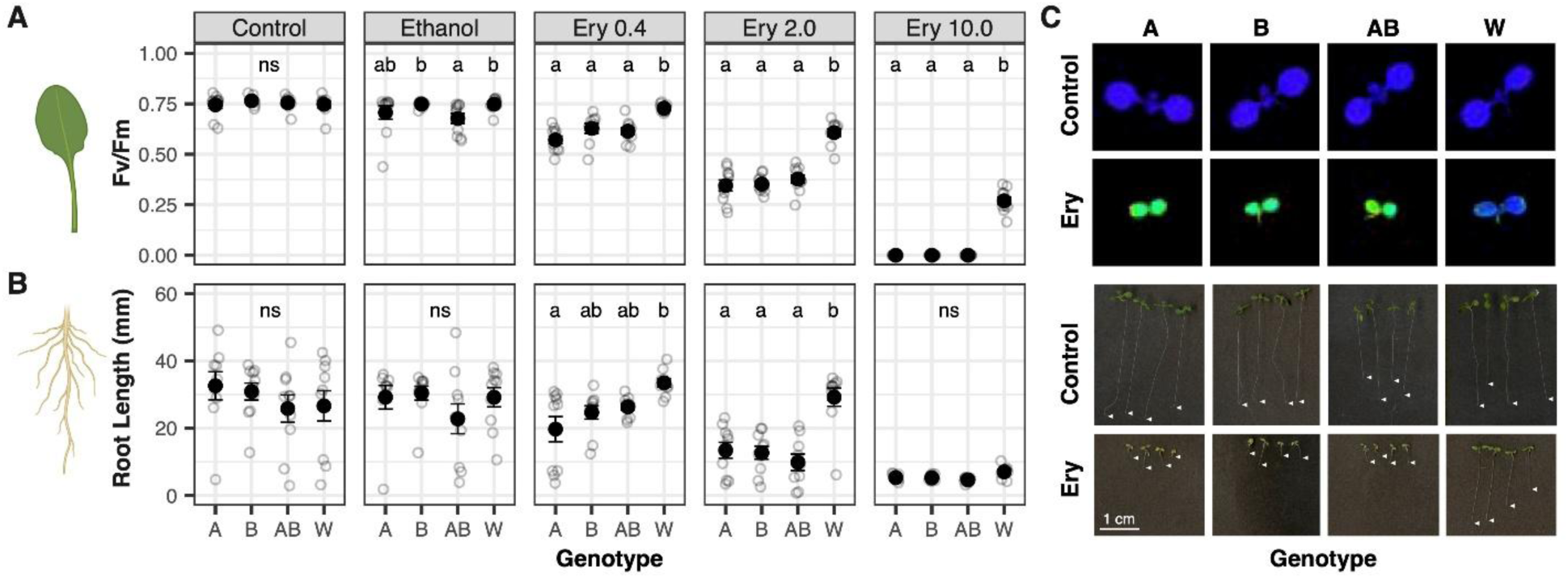
Erythromycin treatment disproportionately impacts photosynthetic efficiency (panel A) and root growth (panel B) of Arabidopsis *mutS2* single and double mutants compared to wild type plants. Seeds of single mutants *mutS2A* (A), *mutS2B* (B), double mutants (AB) and wild type (W) were germinated on media containing increasing concentrations of erythromycin (Ery, concentration in μg/mL). Root length and Fv/Fm measurements were taken after one week of growth (n = 10 individuals of each genotype for each treatment). Plots show mean values as black dots with standard errors. Significant differences (p < 0.05) between genotypes within a treatment are denoted by different letters. Note that the ethanol carrier control showed some variation between genotypes, so the amount of carrier was lowered in subsequent experiments. Panel C shows representative images of chlorophyll fluorescence captured by the PAM fluorometer (top) and root growth of each genotype when exposed to 2 μg/mL Ery. White arrows indicate the root apex of each individual. The leaf and root icons in panels A and B were generated with Biorender.

In bacteria, antibiotics have been identified that bind different areas of the ribosome, stalling them in either rotated or unrotated states. In *B. subtilis*, MutS2 is only able to bind leading ribosomes in the unrotated state due to structural constraints caused by rotation (Cerullo et al. 2022). Thus, at least in *B. subtilis*, *mutS2* mutants are not impacted by antibiotics that lock collided ribosomes in rotated states (Cerullo et al. 2022). To replicate our erythromycin findings and determine whether Arabidopsis MutS2 protein function might also depend on the rotation state of stalled ribosomes, we exposed wt plants and *muts2* single and double mutants to antibiotics that lock ribosomes in either unrotated (erythromycin and chloramphenicol) or rotated states (hygromycin and spectinomycin). We found that both single and double *mutS2* mutants showed disproportionate inhibition in photosynthetic efficiency compared to wt when treated with antibiotics that lock ribosomes in unrotated states (Fig. 3A). Root growth followed a similar trend but was more variable within each genotype (Fig. 3B). By contrast, there was no significant difference among genotypes for either of these phenotypic metrics when plants were treated with antibiotics that lock ribosomes in the rotated state.

**Figure 3.**
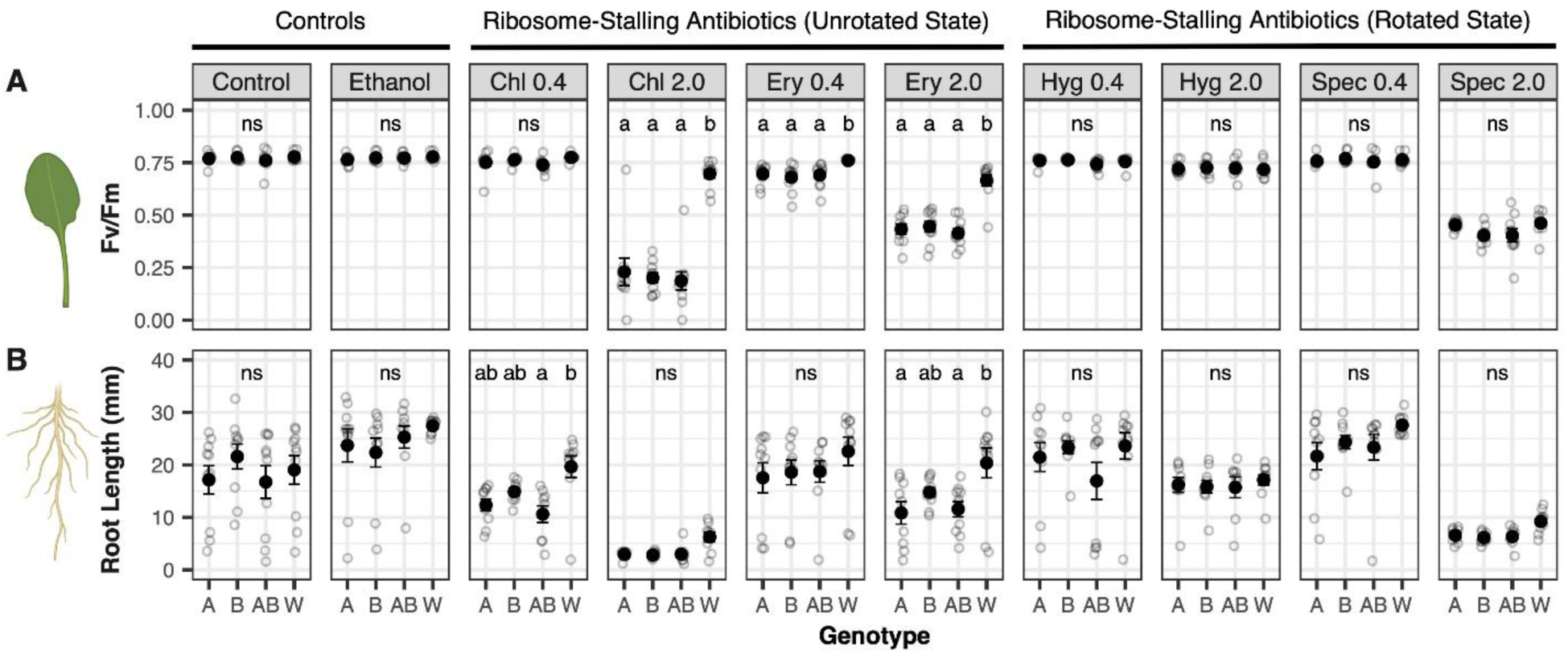
Differential impact of ribosome stalling antibiotics on photosynthetic efficiency (panel A) and root length (panel B) of Arabidopsis *mutS2* mutants. Antibiotics that stall leading ribosomes in unrotated states (chloramphenicol [Chl] and erythromycin [Ery]) disproportionately reduced photosynthetic efficiency of *mutS2* single and double mutants compared to wild type plants. However, treatment with antibiotics stalling ribosomes in rotated states (hygromycin [Hyg] and spectinomycin [Spec]) have similar effects on both wild type and mutant plants. Single mutants *mutS2A* (A), *mutS2B* (B), double mutants (AB) and wild type (W) were grown on plates containing antibiotics for one week before Fv/Fm and root length measurements. Antibiotic concentration is in μg/mL, with two concentrations tested for each antibiotic (n = 10 individuals of each genotype for each treatment). Plots show mean values as black dots with standard errors. Significant differences (p < 0.05) between genotypes within a treatment are denoted by different letters. The leaf and root icons were generated with Biorender.

To confirm that loss of MutS2 function was responsible for the increased sensitivity to erythromycin and chloramphenicol treatment, we complemented *mutS2* single mutants with wt (cDNA) copies of the respective gene via *Agrobacterium*-mediated transformation. We found that reintroducing these genes significantly alleviated the observed Fv/Fm phenotype under erythromycin and chloramphenicol treatment (Fig. 4). The root length data was less clear, and *mutS2A* and double mutant lines exhibited a surprising reduction in growth under control conditions that had not been detected in previous experiments. In general, root growth data showed substantially more variation among replicates than Fv/Fm measures (Figs. 2-4), so this phenotype may be a less reliable assay of MutS2-mediated antibiotic sensitivity than Fv/Fm. Although some individuals in the complementation lines exhibited full restoration to wt levels, only partial rescue was observed in others, especially for *mutS2A* mutants (Fig. 4). Although expression of the reintroduced cDNAs was confirmed by RT-PCR in mature leaf tissue (Supp. Fig. S3), it is possible that the cassettes did not fully mimic native expression of the wt locus in developing seedlings in which the assay was performed. Nevertheless, the observed complementation provides clear evidence that the loss of MutS2 function is responsible for the observed phenotypes.

**Figure 4.**
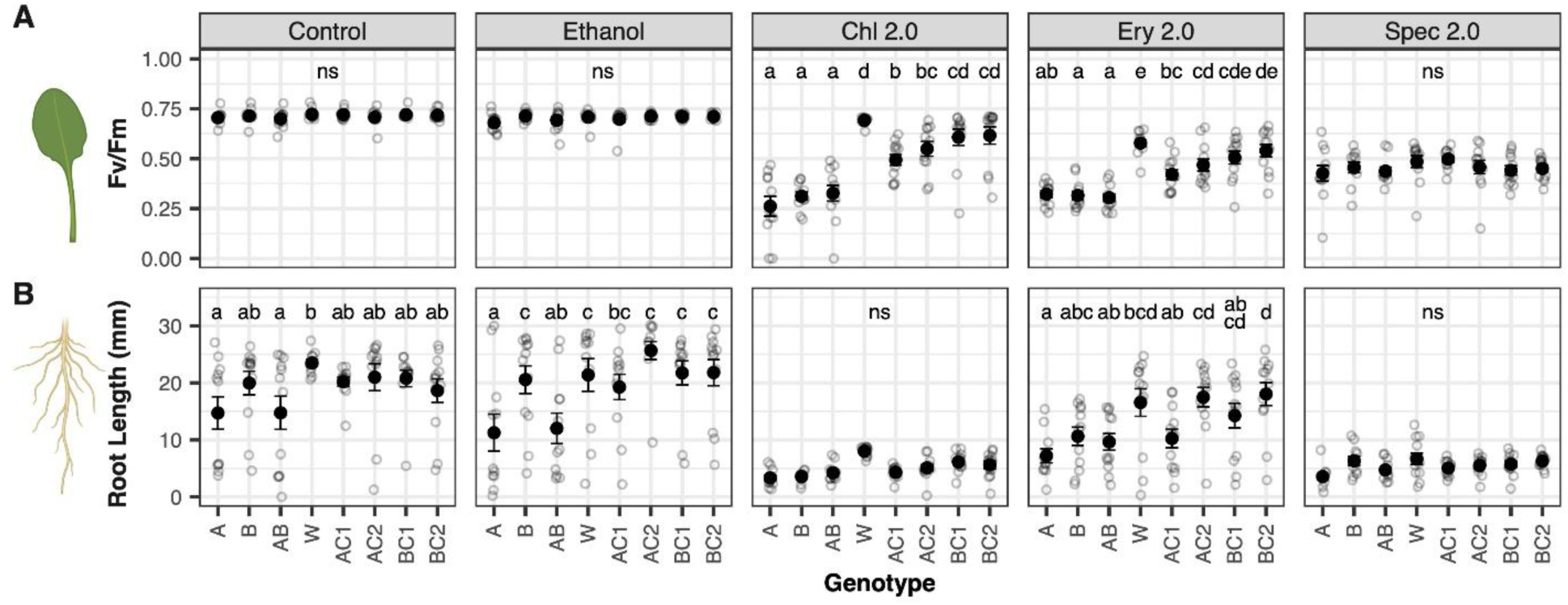
Complementation with transgenic cassettes partially rescue the *mutS2* mutant phenotype. Arabidopsis single mutants *mutS2A* (A) and *mutS2B* (B), double mutants (AB), wild type (W) and lines containing transgenic complementation cassettes (AC1, AC2, BC1, BC2) were grown on plates containing antibiotics for one week before Fv/Fm and root length measurements. Antibiotic concentrations were 2 μg/mL; chloramphenicol (Chl), erythromycin (Ery), and spectinomycin (Spec). Plots show mean values as black dots with standard errors (n = 12 individuals of each genotype for each treatment). Significant differences (p < 0.05) between genotypes within a treatment are denoted by different letters. Hygromycin treatments were not included in this analysis because the complementation cassettes contained a hygromycin resistance selection marker, making any effects of reintroducing *MutS2* genes uninterpretable.

### mutS2 double mutants show small but significant increases in sensitivity to ciprofloxacin and rates of plastid genome rearrangement induced by genotoxic stress

To determine whether MutS2 proteins might also play a role in HR in plastid genomes, we subjected mutant and wt lines to genotoxic stress. Seeds were germinated on media containing the genotoxic agent ciprofloxacin (CIP), an antibiotic that causes double strand breaks (DSBs) in organellar genomes and has been widely used to study mutants in organellar replication, recombination, and repair (Cappadocia et al. 2010; Parent et al. 2011; Miller-Messmer et al. 2012; Schatz-Daas et al. 2022). Treatment with 0.75 μM CIP inhibited the formation of true leaves (i.e., germinated seedlings only produced cotyledons) in *mutS2A/B* double mutants to a greater extent than wt or single mutants (Fig.5). On average, *mutS2B* showed the greatest percentage of plants with true leaves at the two highest CIP concentrations, but this result did not represent a significant difference relative to wt controls (Fig. 5).

**Figure 5.**
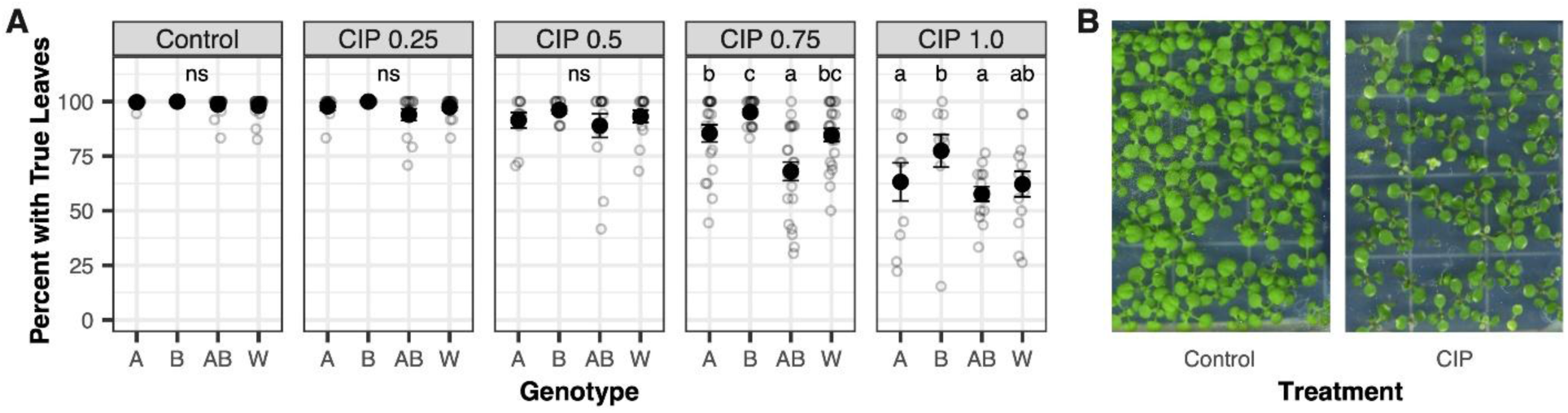
Ciprofloxacin (CIP) treatment impacts development of *muts2A/B* double mutants to a greater extent than single mutants or wild type at 0.75 μM. Arabidopsis single mutants *mutS2A* (A), *mutS2B* (B), double mutants (AB) and wild type (W) seeds were plated on media with increasing concentrations of CIP (concentration shown in μM). The percentage of seedlings that developed true leaves after two weeks of growth is shown in panel A. All other seedlings produced cotyledons only. A small number of seeds did not germinate and were excluded from the analysis. Each data point represents a plate that contained 18 seedlings. Plots show mean values as black dots with standard errors. Significant differences (p < 0.05) between genotypes within a treatment are denoted by different letters. Panel B shows representative images of plants grown in control conditions (left) and 0.75 μM CIP (right).

To further assess the impact of MutS2 on the plastid genome, we extracted DNA from wt and *mutS2A/B* mutant plants grown either with or without CIP (0.75 μM) and performed Illumina sequencing. As expected, we found that treatment with CIP led to extensive organellar genome copy-number variation (Supp. Fig. S4 and S5) and structural rearrangements (Fig. 6A and Supp. Fig. S6) with a repeatable pattern of “hotspots” in the genomes (Supp. Fig. S7). The majority of breakpoint pairs for these rearrangements shared small regions (7-20 bp) of sequence similarity (Fig. 6B), consistent with previous findings indicating that “microhomologies” can be important in the repair of CIP-induced DSBs (Cappadocia et al. 2010; Parent et al. 2011; Miller-Messmer et al. 2012; García-Medel et al. 2019). Under control conditions (no CIP), *mutS2* double mutants did not exhibit any significant difference in the rate of plastid genome rearrangements relative to wt (p = 0.30). By contrast, the impact of CIP treatment was approximately two-fold larger in the mutant background compared to wt (p = 0.0066; Fig. 6A). A similar but only marginally significant trend was found for mitochondrial genome rearrangements, with *mutS2* double mutants showing an approximately 1.7-fold increase compared to wt when treated with CIP (p = 0.0527) (Supp. Fig. S6). This result is surprising given that Arabidopsis MutS2 proteins are only known to be localized to the plastids. Therefore, although these results suggest that MutS2 proteins may play some role in regulating recombination in organellar genomes as observed in bacteria, it remains unclear whether this is a direct effect or a downstream consequence of disrupting another MutS2 function such as RQC.

**Figure 6.**
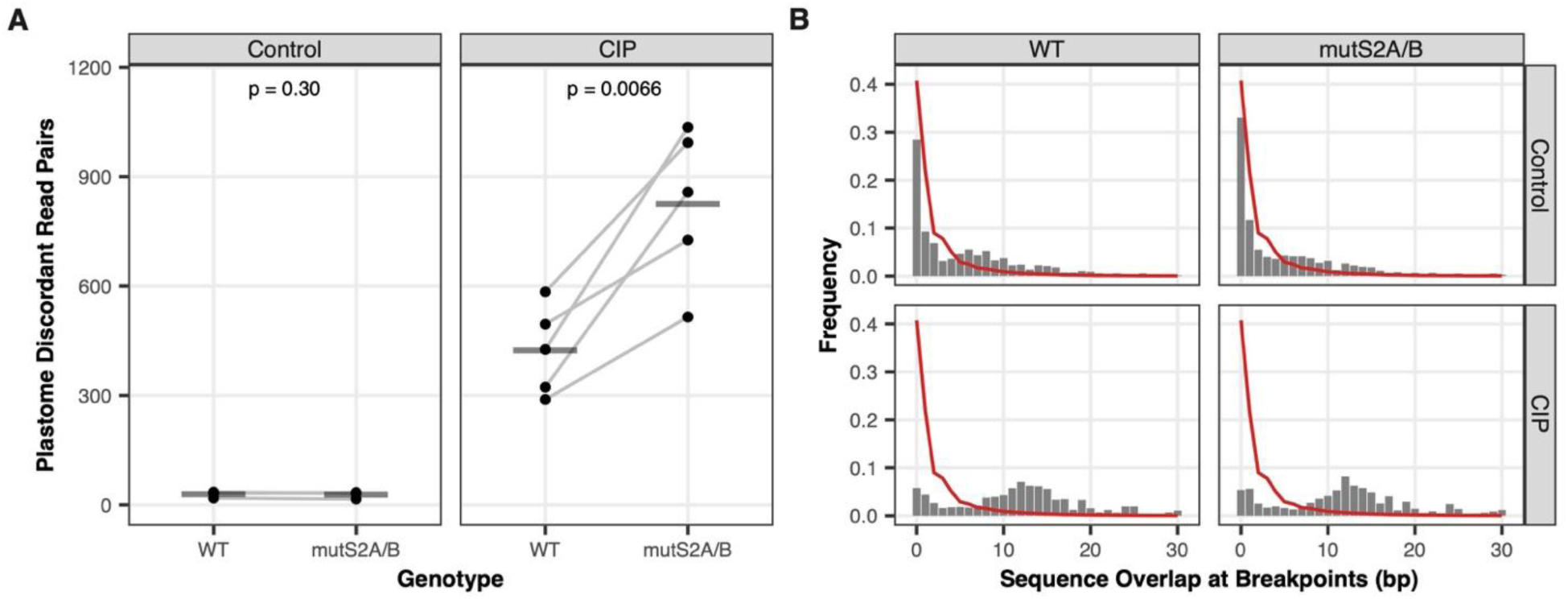
Ciprofloxacin (CIP) disproportionately increases plastid genome (plastome) rearrangements in *mutS2* double mutants compared to wild type. Illumina sequencing was performed on DNA from pools of seedlings treated with or without 0.75 μM CIP, and plastome rearrangements were quantified based on counts of read pairs (per million mapped) that mapped discordantly to the genome with breakpoints positioned within 1 kb of each other. These local rearrangements are characteristic features of plant organellar genome instability (Zampini et al. 2015). (A) Points represent replicate plates containing half wild type and half mutant individuals, with lines connecting data points for samples taken from the same plate. Horizontal bars represent the mean of the five replicates. The reported p-values are based on paired *t*-test comparisons between mutant and wild type within each treatment. (B) The gray bars show the observed frequency distribution for the length of overlapping sequence similarity shared between the pairs of breakpoints for structural rearrangements in the plastid genome. The red lines show the expected distribution based on randomly sampled rearrangement positions.

### De-etiolation response is compromised in mutS2 mutants

Preliminary observations of seedlings under dark treatment suggested that *mutS2* mutants might be inhibited in the process of de-etiolation, which involves greening of cotyledons and development of mature chloroplasts in response to light after etiolation in the dark. To further test for this response, we performed experiments wherein we exposed seeds of wt and *mutS2* single and double mutants to a period of light (8 h) followed by a five-day dark period (resulting in etiolation) and subsequent re-exposure to light. We found that most *mutS2* single and double mutants exhibited a curtailed greening response under these conditions (Fig. 7). The majority of mutant seedlings remained white and undeveloped, even after >2 weeks of light exposure, whereas their wt counterparts turned green and formed true leaves. No differences in greening of the cotyledons were observed among the wt and *muts2* single and double mutant seedlings when grown under control (not subjected to the dark treatment) conditions.

**Figure 7.**
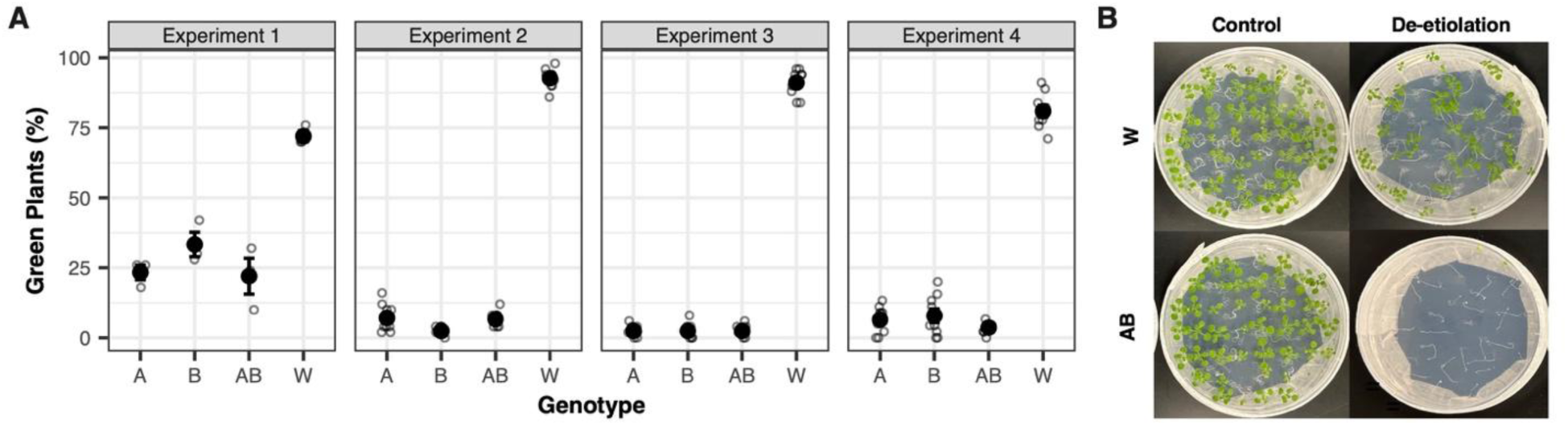
*mutS2* mutants are unable to perform a de-etiolation response. Arabidopsis single mutants *mutS2A* (A) and *mutS2B* (B), double mutants (AB), and wild type (W) seeds were plated on standard MS media, exposed to light to induce germination and then kept in the dark for 5 days. Plants were then re-exposed to light (16/8 hours light/dark), and the number of green seedlings were counted after 1 week. Panel A shows the results of four separate experiments testing the number of de-etiolated plants with green leaves. Open circles represent replicate plates, and mean values are shown as black dots with standard errors. Panel B shows representative examples of plates from the growth of W (top) and AB (bottom) seedlings under normal conditions (not exposed to the etiolation treatment; left) and after etiolation and subsequent re-exposure to light for 2 weeks (right).

## Discussion

### MutS2A and MutS2B are involved in plastid ribosome quality control

Chloroplasts generate extremely high translational demands, yet very little is known about the translational quality control pathways that operate in these organelles. Our work suggests that MutS2A and MutS2B function in plastid RQC, presumably by interacting with plastid ribosomes to resolve collisions. We demonstrate that, like *B. subtilis muts2* mutants, Arabidopsis *muts2* mutants exhibit reduced growth compared to wt when treated with antibiotics that stall ribosomes in unrotated states, but not those that stall ribosomes in rotated states (Fig. 3), suggesting that the mechanism of ribosome stalling and rescue is conserved between plastids and their bacterial relatives. Studies in the green algae *Chlamydomonas reinhardtii* identified MutS2A and MutS2B orthologs (Phytozome protein IDs: Cre01.g040350.t1.1 and Cre01.g014800.t1.1) as plastid ribosome interacting proteins using affinity purification of epitope-tagged plastid ribosomes (Westrich et al. 2021), providing further evidence that these proteins associate with ribosomes.

It is important to note that although antibiotic treatment represents a convenient way to inhibit plastid ribosomes, plant mitochondrial ribosomes might also be sensitive to ribosome-stalling antibiotics, as they too are derived from a bacterial ancestor. Thus, we cannot entirely rule out the possibility that our observed *mutS2* phenotypes may be due to a combination of antibiotic action on both plastid and mitochondrial ribosomes/RQC. The impact of antibiotics on organelle function has gained less attention in plants than animals (Rocha et al. 2021), but chloramphenicol is known to block translation in isolated soybean mitochondria (Mulo et al. 2003). Mammalian mitochondria are known to be negatively impacted by numerous antibiotics, including chloramphenicol and erythromycin, although sensitivity can vary depending on cell line, tissue type and other experimental factors (Duewelhenke et al. 2007; Wang et al. 2015; Xiao et al. 2019; Suárez-Rivero et al. 2021). Unfortunately, antibiotics that specifically target ribosomes of only one organelle have not been characterized, so teasing apart these effects will be challenging. Because empirical studies have not found evidence of mitochondrial localization of MutS2 proteins (Carrie et al. 2009; Fuchs et al. 2020), a role in mitochondrial RQC may be unlikely. Nevertheless, direct tests for physical interactions of MutS2 proteins with mitochondrial and/or plastid ribosomes in Arabidopsis represent an important future direction.

### Potential alternative pathways of ribosome rescue in organelles

Redundant pathways of ribosome rescue are prevalent in bacteria, but it is unclear if this is also the case in plastids. The tmRNA trans-translation system is present in >97% of bacteria (Callan et al. 2024), but it is generally lacking in eukaryotes. However, tmRNA sequences (also known as ssrA) have been identified in plastid genomes of some glaucophytes and red algal lineages, yet they appear to be absent in green algal and land plant plastids (Williams and Bartel 1998; Gueneau de Novoa and Williams 2004). The red alga *Cyanidioschyzon merolae* and the diatom *Thalassiosira pseudonana* contain a nuclear-encoded, plastid-localized tmRNA-binding cofactor SmpB that can function in bacterial trans-translation (Jacob et al. 2005), further suggesting that this system of ribosome rescue may function within some algal plastid genomes. Interestingly, sequence analyses show that, unlike members of the green lineage, red algae and diatoms contain only one full-length copy of MutS2 (Fig. 1) (Berdieva et al. 2024; Sloan et al. 2024). As our results suggest that both MutS2A and MutS2B are required for RQC in Arabidopsis, it remains an open question whether a single MutS2 protein is sufficient for RQC in red algae and diatoms and/or to what extent these lineages might rely on trans-translation versus MutS2-mediated ribosome rescue.

Arf proteins represent a separate type of ribosome rescue that is present in many bacterial taxa (Callan et al. 2024), and homologs of the alternative rescue factor ArfB have been identified in some eukaryotic nuclear genomes. For example, mammalian mitochondria import a nuclear-encoded ArfB homolog (ICT1, Uniprot:Q14197) that confers a lethal phenotype in human cell lines when mutated (Richter et al. 2010). ICT1 can replace the activity of ArfB in bacterial cell lines and can rescue non-stop translation complexes as well as stalled ribosomes with short mRNAs extending past the A site (Feaga et al. 2016). In Arabidopsis, a nuclear-encoded ArfB homolog (At1g62850) was shown to be targeted to plastids and can function in bacterial ribosome rescue assays (Nagao et al. 2020). Phenotypes resulting from the mutation of the plastid ArfB have not yet been characterized, but it will be interesting to see if they parallel that of ICT1 or if this Arf is redundant with other ribosome rescue pathways, similar to its bacterial homologs.

Additional proteins involved in ribosome splitting are known to be important under specific stress conditions. For example, the GTPase known as High Frequency of Lysogenization X (HflX) rescues stalled ribosomes under heat stress in *E. coli* (Zhang et al. 2015). A nuclear-encoded homolog has been identified in human mitochondria (GTPBP6) and shows activity in both ribosome recycling and biogenesis (Lavdovskaia et al. 2020; Hillen et al. 2021). Recently plastid targeted HflX homolog was identified in Arabidopsis (At5g57960) (Mehrez et al. 2024). Although *hflX* mutants showed no significant differences in growth compared to wt under a suite of abiotic stress conditions, they were substantially inhibited when treated with the plastid translation inhibitor lincomycin. Bacterial HflX and its homolog HflxR can overcome lincomycin induced stress either through ribosome splitting (Duval et al. 2018; Rudra et al. 2020) or by binding to the ribosome which results in reduced antibiotic access (Koller et al. 2022). Currently the mode of action and physiological role of Arabidopsis HflX remain unknown.

Redundancy in plastid ribosome rescue pathways could explain the lack of phenotypes in *hflX* and *mutS2* mutants grown under laboratory conditions. However, under specific antibiotic-induced stresses, mutants show distinct phenotypes, suggesting that the products of these genes cannot fully compensate for each other. Similarly, the de-etiolation phenotype of *mutS2* mutants suggests that the presence of ArfB and/or HflX is not sufficient to mitigate this type of translational stress.

Unlike the canonical RQC pathway, Arf and HflX enzymes do not tag proteins for degradation which may lead to a larger disruption in plastid proteostasis. However, it is not yet clear whether the full RQC pathway is present in plastids. Understanding the function of these genes and their potential overlap will provide important insights into how plastids perform ribosome rescue under different developmental and stress conditions.

### Other potential players in plastid RQC

Cleavage of stalled ribosomes by MutS2 may be the first step in the RQC pathway, but other players involved in remedying ribosome collisions in plastids remain entirely uncharacterized. In *B. subtilis*, RqcH (homolog of the eukaryotic Rqc2) functions downstream of MutS2 and encodes an N-fact domain containing protein that promotes degradation of stalled translation products (Lytvynenko et al. 2019; Filbeck et al. 2022). Using BLASTP, we identified a single eukaryotic-like Rqc2 homolog (At5g49930 or EMB1441) in Arabidopsis and no bacterial-like RqcH homologs. Like MutS2A and MutS2B, EMB1441 is widely expressed throughout the plant, but unlike *mutS2* mutants, which show no phenotype under normal growth conditions, *emb1441* mutants are embryo lethal (Rhee et al. 2003; Waese et al. 2017; Meinke 2020). Given the ancestral eukaryotic origins of this Rqc2 homolog, it likely functions in cytosolic RQC, which may explain its essential role. Although subcellular localization databases have not reported it as plastid-localized (Forsythe et al. 2019), a small number of proteomic studies have identified EMB1441 peptides in chloroplasts, including thylakoid membrane fractions harvested from seedlings undergoing de-etiolation (van Wijk et al. 2021). Therefore, it is possible that this sole homolog of RqcH/Rqc2 in Arabidopsis has been recruited to function in plastid (bacterial-like) RQC. Future studies should identify the localization and binding partners of Arabidopsis EMB1441 to determine whether it has a dual function in both plastid and nuclear RQC. If this protein is not targeted to plastids, it may indicate that an entirely different pathway operates downstream of MutS2. For example, it is possible MutS2 proteins serve to initiate ribosome recycling but the aborted polypeptides released from stalled ribosomes are degraded through more general plastid proteolysis pathways (van Wijk 2015, 2024; Nishimura et al. 2017).

RqcP, which functions downstream of RqcH in most bacteria that utilize RQC, is lacking in eukaryotic RQC pathways and also appears to be absent from Arabidopsis based on homology search. The final step in the RQC pathway targets nascent polypeptides for degradation, either through alanine (Ala) tailing (bacterial) or ubiquitination (eukaryotic) (Filbeck et al. 2022). In *B. subtilis*, RqcH is responsible for Ala tailing, which targets nascent polypeptides to the ClpXP protease (Lytvynenko et al. 2019; Crowe-McAuliffe et al. 2021; Filbeck et al. 2021; Takada et al. 2021). However, in the eukaryotic system, Rqc2 adds Ala-Thr tails and recruits a member of the E3 ubiquitin-ligase pathway (Ltn/Listerin) to target polypeptides for degradation (Shen et al. 2015; Doamekpor et al. 2016). Currently, there is no conclusive evidence that ubiquitin or E3 ligases exist within the plastid (van Wijk et al. 2023). However, very little is known about the specific features of most plastid degradation signals, although they are presumed to be similar to their ancestral bacterial counterparts (van Wijk 2024). How the plastid recognizes and degrades aberrant peptides that result from ribosome stalling remains to be seen.

### A potential role for MutS2 in plastid genome recombination

Although MutS2 proteins are classically known for their role in HR (Eisen 1998; Sachadyn 2010), we only found weak evidence to support this function in Arabidopsis. Our *muts2A/B* double mutants showed slightly increased sensitivity to CIP-induced organellar DSBs (Fig. 5), and they exhibited small (∼2-fold) increases in the rate of CIP-induced rearrangements in the plastid genome compared to wt (Fig. 6A). These results suggest that plant MutS2 proteins may be another of the many factors that mediate recombination and DSB repair in plastid genomes (Maréchal and Brisson 2010; Zhang et al. 2016). However, this interpretation is complicated by the small magnitude of the observed effects and the fact that CIP-treated *muts2A/B* double mutants also showed a trend towards increased rearrangements in mitochondrial genomes compared to wt (Supp. Fig. S6). Therefore, we propose two alternative explanations. First, as discussed in relation to RQC, it is possible that MutS2 proteins function in recombination pathways in both plastids and mitochondria. Alternatively, the observed effects on plastid and/or mitochondrial genome stability may be indirect consequences of the cellular stress arising from a combination of CIP treatment and the loss of plastid RQC function. Testing plant MutS2 binding affinity for various DNA substrates (including recombination intermediates) is a natural next step for distinguishing between these two alternative explanations.

### Protein structure and the functional relationship between MutS2A and MutS2B

The conservation of both copies of MutS2 across Viridiplantae (Fig. 1) suggests that they provide important and distinct functions. In our assays using CIP to induce DSBs in organelle genomes (Fig. 5), we saw adverse phenotypes in double mutants but not in single mutants, suggesting MutS2A and MutS2B may act redundantly in this role. However, we found that MutS2A and MutS2B are both required to mitigate effects induced by the ribosome stalling antibiotics erythromycin and chloramphenicol, suggesting they serve non-redundant functions in RQC (Fig. 3). Thus, the ancient conservation of both gene copies may be due to their role in remedying ribosome collisions.

Bacterial MutS2 functions as a homodimer, and dimerization of the two proteins forms the clamp like structure that is required for binding of both DNA (Fukui et al. 2022) and the leading stalled ribosome (Cerullo et al. 2022). Because both MutS2A and MutS2B appear to be required for RQC, we speculate that these two proteins function as a heterodimer, but future studies are needed to test this prediction. Other instances of heterodimerization are known in the broader MutS family. For example, heterodimerization of human MSH4 and MSH5 is required for their function in homologous recombination in germline cells (Snowden et al. 2004). In any case, dimerization occurs via the Walker A/B type ATPase domain, which is highly conserved across the MutS family (Sachadyn 2010; Ogata et al. 2011) and is present in both plant MutS2A and MutS2B.

The main structural difference between plant MutS2A and MutS2B is the absence of the Smr endonuclease domain in MutS2A (Fig. 1). In *E. coli*, which lacks MutS2 entirely, SmrB is critical in resolving ribosome collisions and performs this act by cleaving the mRNA between the stalled and collided ribosomes (Saito et al. 2022). Similarly, the Smr domain containing protein Cue2 from yeast is recruited to stalled ribosomes and cleaves mRNAs, targeting them for decay (D’Orazio et al. 2019). Recent work has shown that although the *B. subtilis* MutS2 Smr domain enhances binding of MutS2 to collided ribosomes, it is not involved in mRNA cleavage (Park et al. 2024). The catalytic residues of Smr domains in various proteins have been identified, but there is uncertainty in the literature about which specific residues are critical for function (Fukui et al. 2008; Fukui and Kuramitsu 2011; Damke et al. 2015; Zhou et al. 2017; D’Orazio et al. 2019; Saito et al. 2022; Sloan et al. 2024). However, these sites could differ based on the protein’s affinity for and activity against RNA versus DNA. Additional studies will be required to determine whether Arabidopsis MutS2B and its Smr domain have nuclease activity with either of these substates.

### The role of MutS2 during the de-etiolation response

Both *mutS2* single and double mutants showed extreme inhibition during the de-etiolation response and were, for the most part, unable to produce green tissue after re-exposure to light (Fig. 7). We hypothesize that de-etiolation may act as a stressor that activates MutS2 and the RQC pathway, as intense reprogramming occurs in the plastid during the transition from etiolated tissue to green tissue (Jarvis and López-Juez 2013; Pipitone et al. 2021). Previous work has shown that the number of ribosomes occupying an mRNA is the main driver of tissue greening (Legen and Schmitz-Linneweber 2017; Armarego-Marriott et al. 2020). Thus, the transition from etioplasts to mature chloroplasts requires a high translational demand that could increase pressure on RQC due to amino acid deficiencies and/or simply the abundance of mRNAs being co-translated, both of which would create more opportunities for ribosome collisions. A delayed de-etiolation response is also observed in mutants lacking the plastid nucleoid protein NusG, which is involved in coupling chloroplast transcription and translation (Xiong et al. 2022), further connecting a reduction in translational efficiency with this phenotype.

Additionally, as etioplasts (the non-pigmented plastids in etiolated tissues) transition to mature chloroplasts, photosynthetic reactions occur and begin to generate ROS. If left unchecked, these ROS may damage both RNA and DNA in the cell. Not only could this increase ribosome collisions by creating aberrant mRNAs, it also has the potential to damage plastid DNA. Interestingly, MutS2 is upregulated during oxidative stress in *H. pylori* and *D. radiodurans* (Wang et al. 2005; Zhang et al. 2014), and *H. pylori* MutS2 has been shown to bind and repair 8-oxo-G which is a frequent by-product of oxidative stress (Wang and Maier 2017). Thus, the potential roles of MutS2 proteins in both RQC *and* HR may be an important piece of the de-etiolation response, as rapid replication of plastid DNA is required for organelle division and subsequent plant growth. Regardless of the mechanism, the de-etiolation phenotype highlights the importance of MutS2A and MutS2B function in plant growth and development. More clearly understanding the specific role of these proteins in the cell and the mechanisms behind plastid RQC in general will shed light on this neglected area of study.

## Methods

### Phylogenetic, domain architecture, and in silico subcellular targeting analyses

MutS2 protein sequences were obtained from a diverse set of land plants, green algae, red algae, glaucophytes, and bacteria (Supp. Fig. S1, sequences at https://github.com/dbsloan/plant_mutS2_phenotypes) and aligned with MAFFT v7.487 (Katoh and Standley 2013) under default settings. The resulting alignments were trimmed with GBLOCKS v0.91.1 (Castresana 2000) under default settings followed by manual removal of unalignable sequences. Maximum-likelihood phylogenetic inference was performed with RAxML v8.2.12 (Stamatakis 2014), using a PROTGAMMALG model with 100 starting trees and the rapid bootstrap option (-f a). Mitochondrial and plastid targeting predictions were performed with three different in silico methods, using “plant” settings in each case: LOCALIZER, TargetP 2.0, and Predotar v1.04 (Small et al. 2004; Sperschneider et al. 2017; Almagro Armenteros et al. 2019). Domain architecture was predicted with the NCBI CD Search Tool (Conserved Domain Database v.3.19) (Yang et al. 2020). Visualization of the phylogenetic tree, targeting data, and domain architectures was performed with custom scripts using the R ggtree package (Yu 2020).

### Coexpression and evolutionary rate covariation analysis

For coexpression analysis, *MutS2A* (At5g54090) and *MutS2B* (At1g65070) were used as queries in the ATTED-II database (https://atted.jp/) (Obayashi et al. 2022). First, the coexpression viewer was used to perform a pairwise analysis between MutS2A and MutS2B. Then a list of the top 300 coexpressed genes was generated individually for MutS2A and MutS2B using the Arabidopsis ‘coex’ function (Supp. Table S1 and Supp. Table S2). Each gene list was submitted to Panther for GO analysis (http://www.pantherdb.org/) (Thomas et al. 2022), using the statistical overrepresentation test (Mi et al. 2019) with the Arabidopsis database as a reference gene set (Supp. Table S3). GO lists for both genes were sorted for overrepresentation of terms (underrepresented terms were not used in the analysis). GO terms appeared highly enriched for certain broad categories, so we manually curated these terms to place them into general groups including DNA, RNA, DNA/RNA (‘nucleic acid’ in GO term), ATPase, translation (including ‘tRNA’ in GO term), chloroplast, and development.

We also used both *MutS2A* and *MutS2B* as queries to search for signatures of ERC against a previously compiled dataset of 7938 genes from 20 angiosperms (Forsythe et al. 2021), using a phylogenomic pipeline available at: https://github.com/EvanForsythe/Plastid_nuclear_ERC. To determine statistical significance, we evaluated p-values from both Pearson and Spearman correlation coefficients, which we corrected for multiple tests using false discovery rate (Benjamini and Hochberg 1995). ERC hits were designated as genes that display both a Pearson and Spearman uncorrected p-value less than 0.01 and an R^2^ value greater than 0.5 (Supp. Table S4).

### Mutants and crossing design

Mutant lines of *mutS2A* (At5g54090; CS878469) and *mutS2B* (At1g65070; SALK_206291C) were obtained from Arabidopsis Biological Resource Center (Rhee et al. 2003). Both lines contain T-DNA insertions in the first exon of the coding sequence and are in the *Arabidopsis thaliana* Columbia-0 background. Seeds were stratified at 4 °C in water for 3 days and then sown directly into ProMix BX media and grown on shelves with fluorescent light (∼100 µE m^-2^ sec^-1^) at ∼22-23 °C under long-day conditions (16 hr light / 8 hr dark). Homozygous mutant plants were identified by genotyping (primers listed in Supp. Table S5) and reciprocally crossed to generate double heterozygotes, which were subsequently self-pollinated to obtain homozygous double mutants (*mutS2A/B*). To obtain a segregating population for experimentation derived from a wt (“clean”) cytoplasmic background, wt females were crossed with homozygous double mutants (*mutS2A/B*) as males. The resulting heterozygous F1 was self-pollinated, and segregating single mutant, double mutant, and wt F2 plants were identified by genotyping (Supp. Table S5). Seeds of these plants were collected for subsequent experiments. For genotyping, genomic DNA was extracted from leaves by grinding tissue in 200 mM Tris-HCl pH 9, 0.25 M ethylenediaminetetraacetic acid (EDTA) and 1% sodium dodecyl sulfate (SDS) followed by precipitation in isopropanol.

### Complementation lines

MutS2 complementation cassettes were created by cloning the full-length cDNA of either *MutS2A* (At5g54090.1) or *MutS2B* (At1g65070.2) in the binary vector pCambia 1390. Primers are listed in Supp. Table S5, and plasmids have been deposited to Addgene (accessions 246544 and 246545). Constructs were confirmed by sequencing and transformed into *Agrobacterium tumefaciens* GV3101. Plants were transformed by floral dip (Zhang et al. 2006) and selected on hygromycin using previously described methods (Harrison et al. 2006).

### Reverse transcriptase PCR

Approximately 100 mg of leaf tissue was harvested, frozen and disrupted using the TissueLyser (Qiagen). RNA was extracted using the Qiagen RNeasy Plant Mini Kit. RNA was quantified using a Qubit, and 700 ng was treated with DNase (Thermo Scientific EN0521) following the manufacturer’s instructions. cDNA was synthesized from 500 ng of DNase-treated RNA using BioRad iScript RT Supermix. Controls using the –RT Supermix were also performed for each sample. Expression was examined using PCR consisting of 0.5 µL cDNA, 1x Paq5000 Buffer, 0.8 µl of 100 mM dNTPs, 0.5 µl of each 5 µM primer (Supp. Table S5) and 0.1 µL of Paq5000 (Agilent) in a total volume of 10 µL. A gDNA sample (and plasmid DNA for complementation assays) was also used as a template for a positive control. Thermal cycling was performed with a 3 min 95 °C hot start followed by 35 cycles of 95 °C 30 sec, 55 °C 30 sec, 72° C 40 sec, and a final extension at 72 °C for 5 min.

### Plant media and growth conditions

For experiments done on plates, seeds were sterilized in 50% bleach + 1 drop tween for 10 min, washed a minimum of 3 times in sterile water and placed onto square plates using a pipette tip.

Plates were sealed with Parafilm, and seeds were stratified at 4° C for 3 days before being placed in a growth chamber (light intensity 50 µE m^-2^ sec^-1^) at 23° C. Standard media consisted of ½ strength MS containing Gamborg’s vitamins (Cassion MSP06), 0.5 g/L MES (pH to 5.7 with KOH), 1% sucrose (unless otherwise noted), and 0.8% phytoagar (Cassion PTP01). Approximately 50 mL of media was poured into each 100 mm × 100 mm gridded square plate (36 squares in each grid).

### Treatment with ribosome-stalling antibiotics

To identify potential effects on ribosome collisions, antibiotics (erythromycin (Sigma E5389), chloramphenicol (OmniPur 3130), hygromycin (Gold Biol H-270), and spectinomycin (Gold Bio S-140)) were resuspended in 50% ethanol (stock concentration of 25 mg/mL) and added to standard media lacking sucrose at a final concentration of 0, 0.4, 2.0, or 10 µg/mL. The carrier (50% ethanol) was also included as a control treatment because initial experiments showed that ethanol can inhibit the growth of *mutS2* mutants. In initial experiments, two lines of each genotype (wt, *muts2A*, *muts2B*, *muts2A/B*) were sown at the top of a square gridded plate in random order (eight seeds per plate) with six replicates per treatment. Subsequent experiments were performed in a similar manner, but only two antibiotic concentrations were used (0.4 and 2.0 µg/mL) in addition to the control, with five replicate plates per treatment. For experiments with complementation cassettes, one line of each genotype and two lines of each T2 complementation cassette were sown on plates containing 2.0 µg/mL of antibiotic, and this was replicated 12 times for each treatment. For all experiments, plates were placed horizontally in random order in a growth chamber, and plants were grown for 1 week. Plants were dark adapted for at least 20 minutes and subjected to analysis using a PAM Fluorometer (WALZ IMAG-K6) to obtain Fv/Fm readings. In addition, images of all plates were obtained using a scanner and root lengths were measured using ImageJ v1.54. Data were analyzed using a liner mixed effects model in R (lme4 package) with genotype, drug condition, and genotype*drug condition as fixed effects and plate, randomized plate location, and plant line as random effects.

### Ciprofloxacin treatment and analysis of plastid genome rearrangements

To test homologous recombination phenotypes, we used CIP (Thermo Scientific J6137.06), a quinoline antibiotic that creates DSBs in plant organellar genomes. CIP was resuspended in 0.1M HCl, diluted in water and added to media at a final concentration of either 0, 0.25, 0.5, 0.75 or 1.0 µM (as in (Schatz-Daas et al. 2022)). Plates were divided into four equal quadrants, and 18 seeds of a single line of wt, *mutS2A*, *mutS2B*, and *mutS2A/B* were placed into separate quadrants on each plate. Four or more replicate plates were made for each concentration of CIP. After two weeks of growth, counts were made to determine the proportion of plants that had developed true leaves.

The number of ungerminated seeds was low for all lines, and these were not included in the analyzed count data. The experiment was repeated three additional times using the same lines/genotypes, and data were analyzed in R (lme4 package) using a mixed linear model with genotype, CIP concentration, and the interaction (genotype*CIP concentration) as main effects, and with experiment and plate as random effects.

In the final CIP experiment, whole plants of each genotype within a plate (∼18 plants) were pooled and flash frozen in liquid nitrogen. Paired samples of wt and *mutS2A/B* double mutants grown on either 0 or 0.75 µM CIP were chosen for DNA extraction, with five replicates of each. Tissue was pulverized using a TissueLyser (Qiagen), DNA was isolated with the DNeasy Plant Mini Kit (Qiagen), and DNA was concentrated using the Zymo DNA Clean and Concentrator-5 kit (D4013). Illumina libraries were prepared and sequenced by Novogene, using an NEB Ultra II Kit and a 2×150 run on the NovaSeq 6000 platform. The resulting sequencing reads were trimmed with Cutapadt v4.0 (Martin 2011) to remove adapters and low-quality sequence, using a maximum error rate of 0.15, a base quality threshold of 20, and a minimum read length of 75. The first 45 bp of each read in pair were then mapped to the *A. thaliana* Col-0 plastid reference genome (GenBank NC_000932.1) with the second copy of the inverted repeat removed and mitochondrial reference genome (NC_037304.1), using bowtie2 v2.2.5 (Langmead and Salzberg 2012). The maximum length for concordant read mapping was set to 1000 bp (-X 1000 parameter in bowtie2). Total read depth, as well as discordant read pair counts, positions, and sequence overlap at break points were extracted and analyzed, using custom scripts available via https://github.com/dbsloan/plant_mutS2_phenotypes.

## Data and Code Availability

Illumina sequencing reads were deposited into the NCBI Sequence Read Archive (BioProject PRJNA1298224). All phenotypic datasets and scripts for statistical analysis and figure generated are available via GitHub (https://github.com/dbsloan/plant_mutS2_phenotypes).

## Supporting information

All supplemental tables

## Acknowledgements

This research was supported by grants and fellowships from the National Institutes of Health (R35GM148134 to DBS and T32GM132057 to SAK) and the National Science Foundation (IOS-2114641 to DBS and ESF and GRFP-006784 to SAK).

## Author contributions

AKB designed and performed research, analyzed data, wrote manuscript. KK performed research, analyzed data. KVSKP designed and performed research, analyzed data. SAK designed and performed research, analyzed data. SL performed research, analyzed data. EFS performed research, analyzed data. DBS designed and performed research, analyzed data, wrote manuscript. All authors revised the manuscript and approved the final version.

**Supplemental Figure S1.**
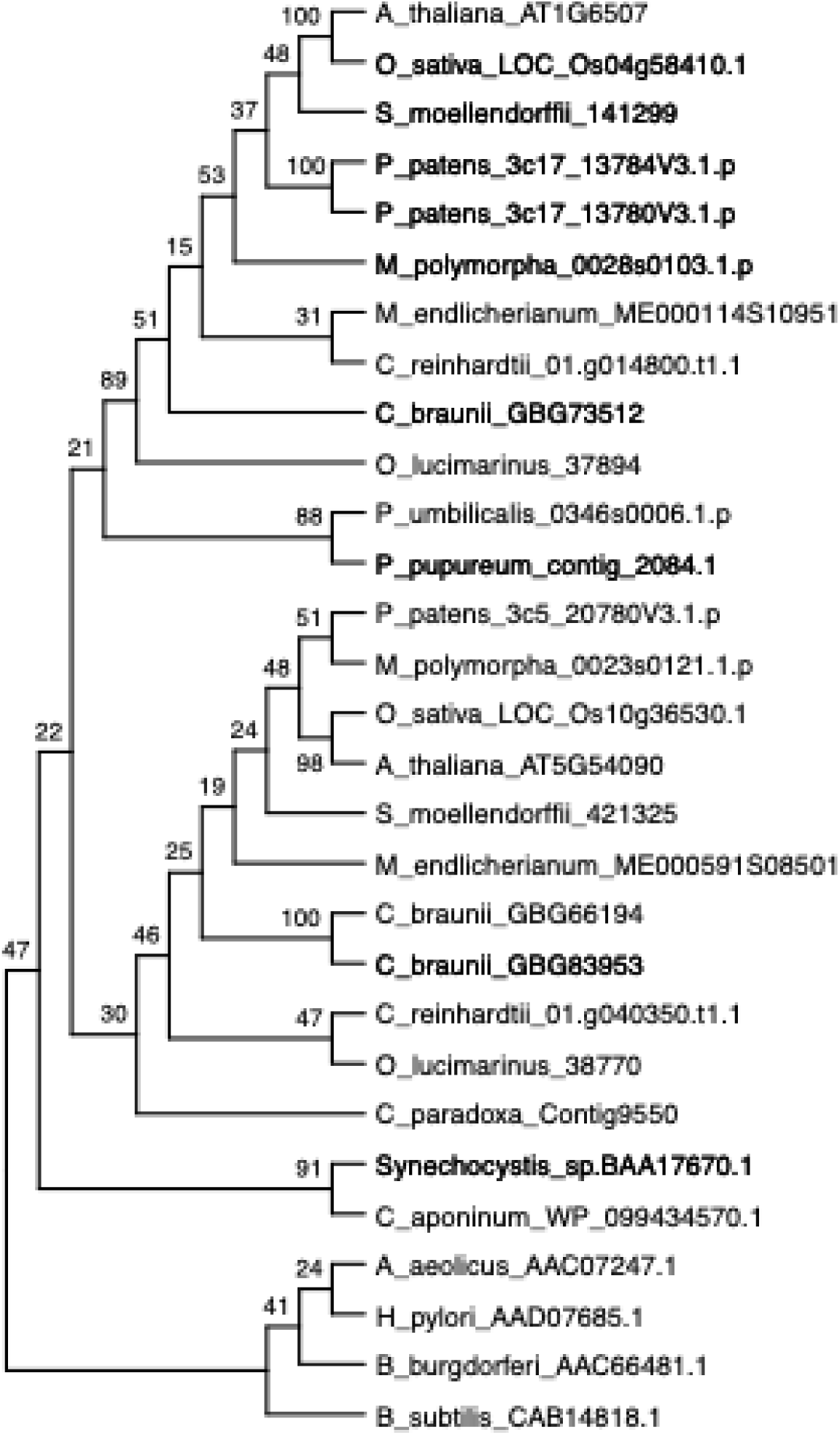
Maximum-likelihood tree inferred from MutS2 protein sequences. This tree corresponds to the one presented in Fig. 1, but bootstrap support values and sequence accessions are indicated. Note that the tree is ultrametric (i.e., branch lengths are not scaled to sequence divergence).

**Supplemental Figure S2.**
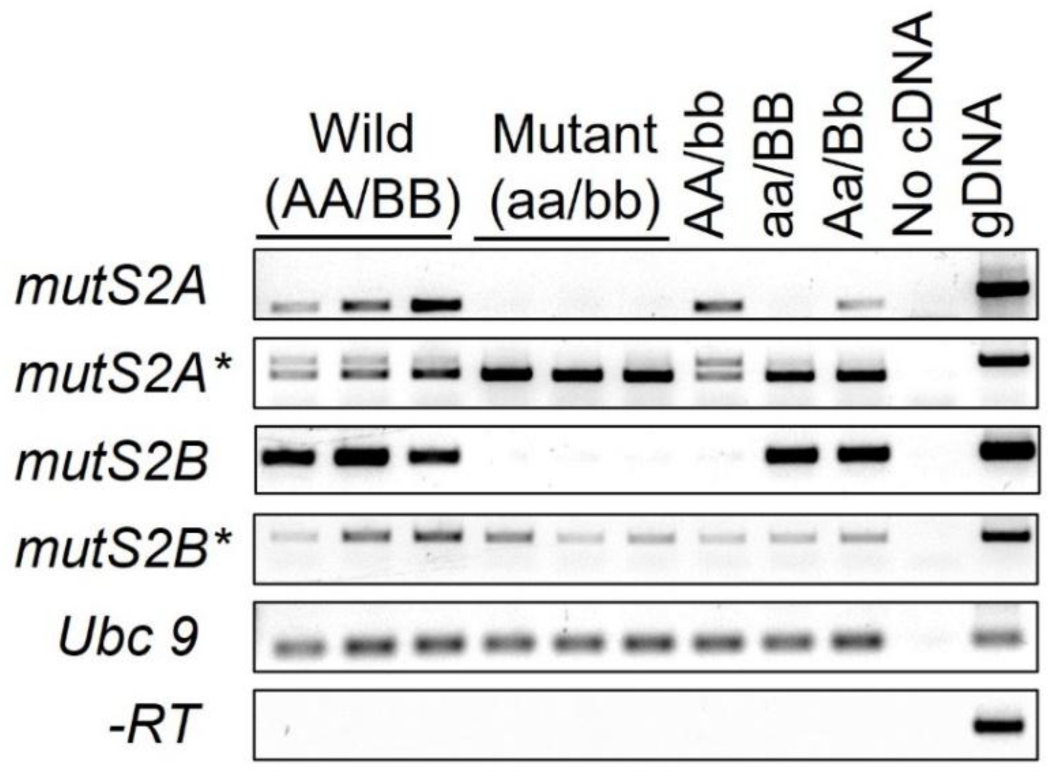
Reverse transcriptase PCR of *MutS2* alleles in plants from a segregating F2 population. Plant genotypes are listed along the upper part of the plot and primers are listed on the left-hand side. The T-DNA insertion for both MutS2A and MutS2B lines is located in the first exon and is expected to disrupt translation of this exon and the signal peptide. Primers spanning the T-DNA insertion (*mutS2A*, *mutS2B*) region show no amplification in homozygous mutants as expected. However, primers downstream of the insertion (*mutS2A*, mutS2B\**) show amplification in all samples. Primer set *Ubc9* was used as a positive control. *-RT* indicates no reverse transcriptase was added during cDNA amplification. Plant genotypes: Wild (AA/BB), homozygous wild type at both loci; Mutant (aa/bb), homozygous mutant at both loci; AA/bb, homozygous wild type at *MutS2A* locus, homozygous mutant at *MutS2B* locus; aa/BB, homozygous mutant at *MutS2A* locus, homozygous wild type at *MutS2B* locus; Aa/Bb, heterozygous at both loci; No cDNA, no template added; gDNA, genomic DNA from heterozygous (F1) parent plant.

**Supplemental Figure S3.**
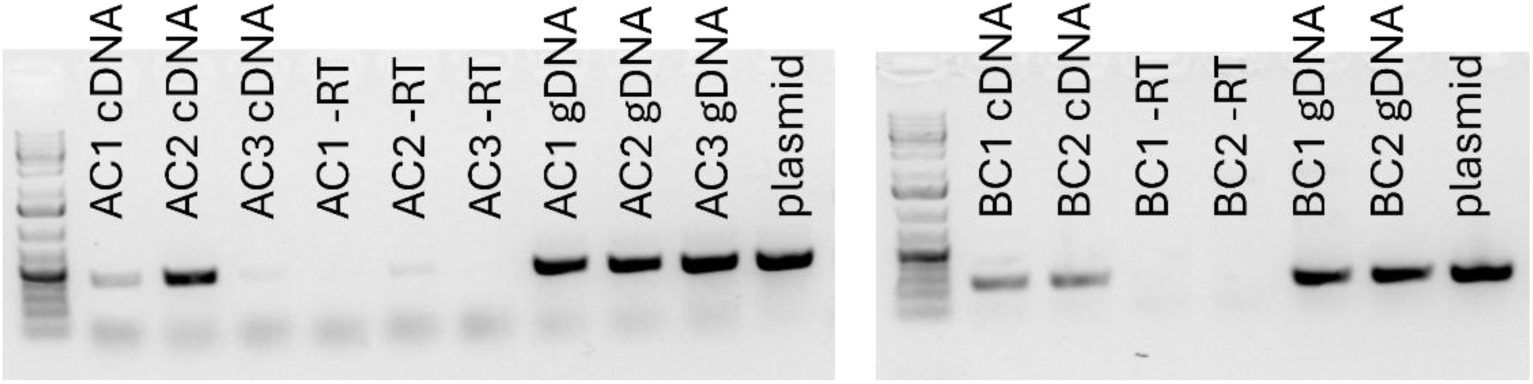
Reverse transcriptase PCR of re-introduced *MutS2* cDNAs in plants containing complementation cassettes. AC and BC refer to MutS2A and MutS2B complementation lines, respectively. –RT indicates no reverse transcriptase was added during cDNA amplification. Primers spanning the T-DNA insertion region were used for RT-PCR as these showed no expression in muts2 mutants (Supp. Fig. S2)

**Supplemental Figure S4.**
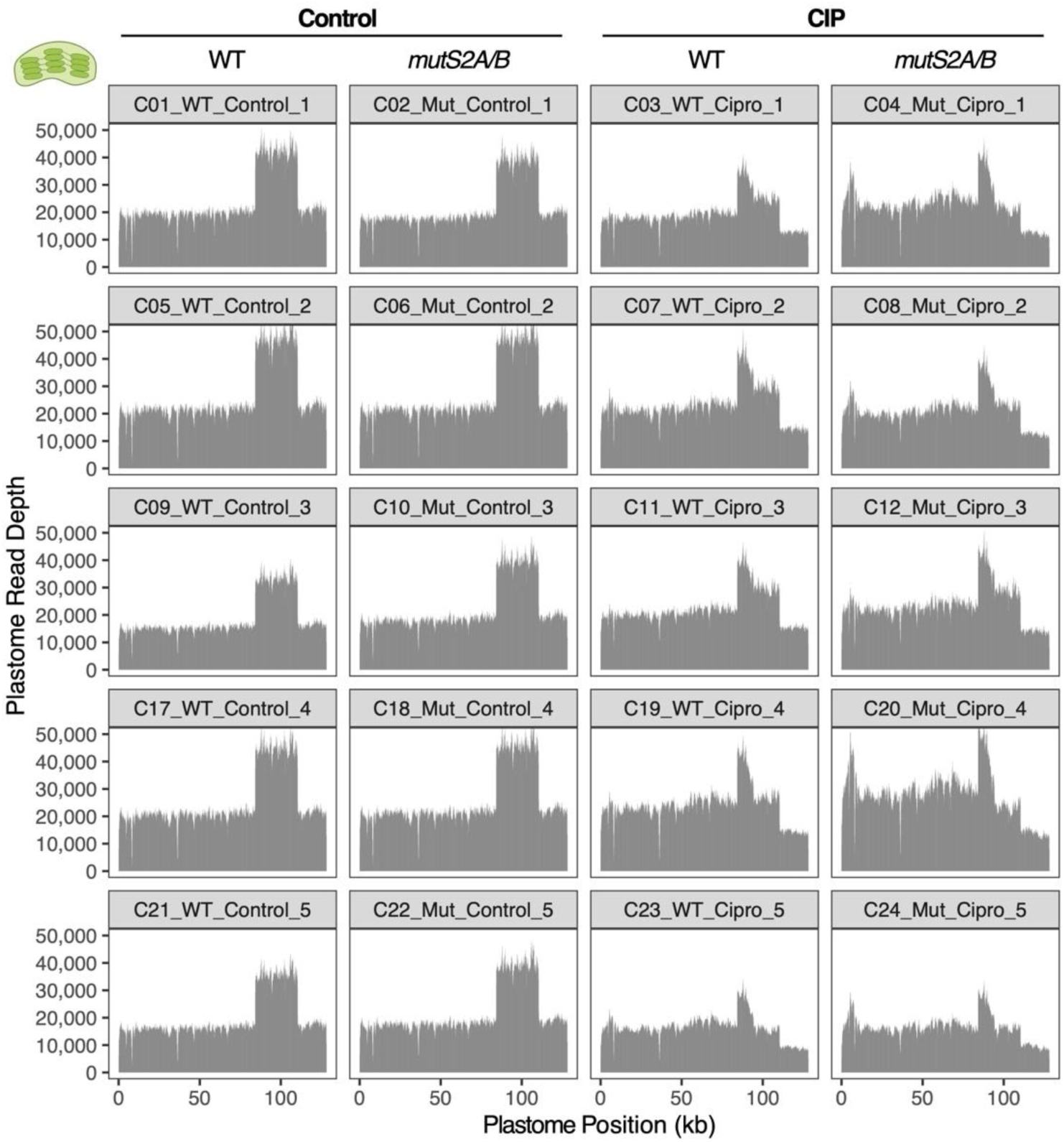
Plastid genome (plastome) sequencing coverage in Arabidopsis grown under control conditions (left) or treated with 0.75 μM ciprofloxacin (CIP, right). Each treatment was applied to wild type plants (first and third columns) and *mutS2* double mutants (second and fourth columns). Five biological replicates were included for each treatment/genotype combination. The x-axis indicates position along the length of the Arabidopsis plastome, and the y-axis indicates the depth of sequencing coverage in 100-bp sliding windows. The second copy of the inverted repeat was removed from the reference for mapping purposes, which explains why all control samples have approximately two-fold higher coverage across the first copy of the repeat. Note that the CIP-treated samples consistently show drops in coverage in part of the inverted repeat and small single-copy region, indicative of a repeatable pattern of structural rearrangements.

**Supplemental Figure S5.**
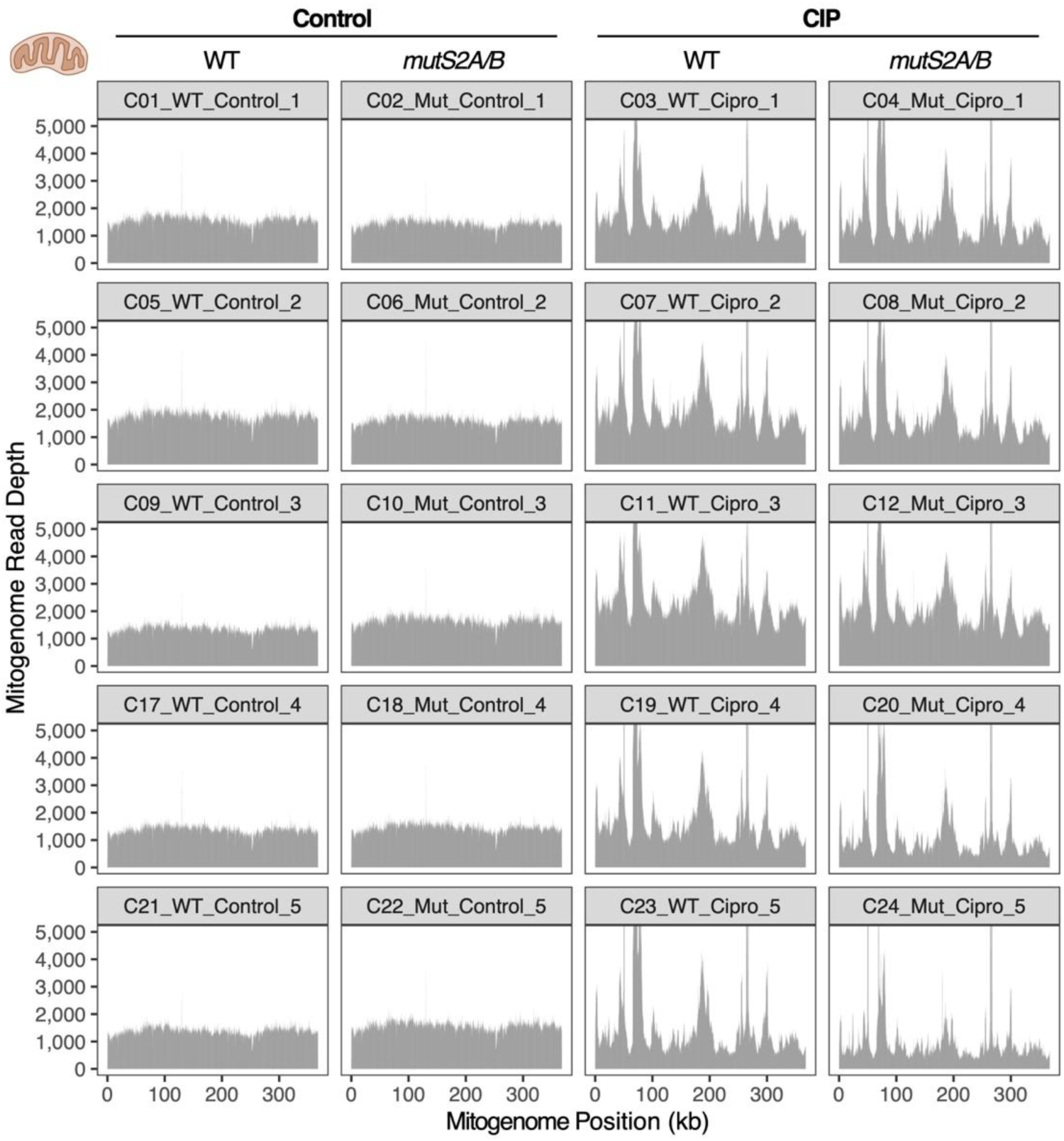
Mitochondrial genome (mitogenome) sequencing coverage in Arabidopsis grown under control conditions (left) or treated with 0.75 μM ciprofloxacin (CIP, right). Each treatment was applied to wild type plants (first and third columns) and *mutS2* double mutants (second and fourth columns). Five biological replicates were included for each treatment/genotype combination. The x-axis indicates position along the length of the Arabidopsis mitogenome, and the y-axis indicates the depth of sequencing coverage in 100-bp sliding windows. Note that the CIP-treated samples exhibit a pattern of extreme variation in sequencing depth across the genome that is highly consistent across replicates and genotypes, indicative of a repeatable pattern of structural rearrangements.

**Supplemental Figure S56.**
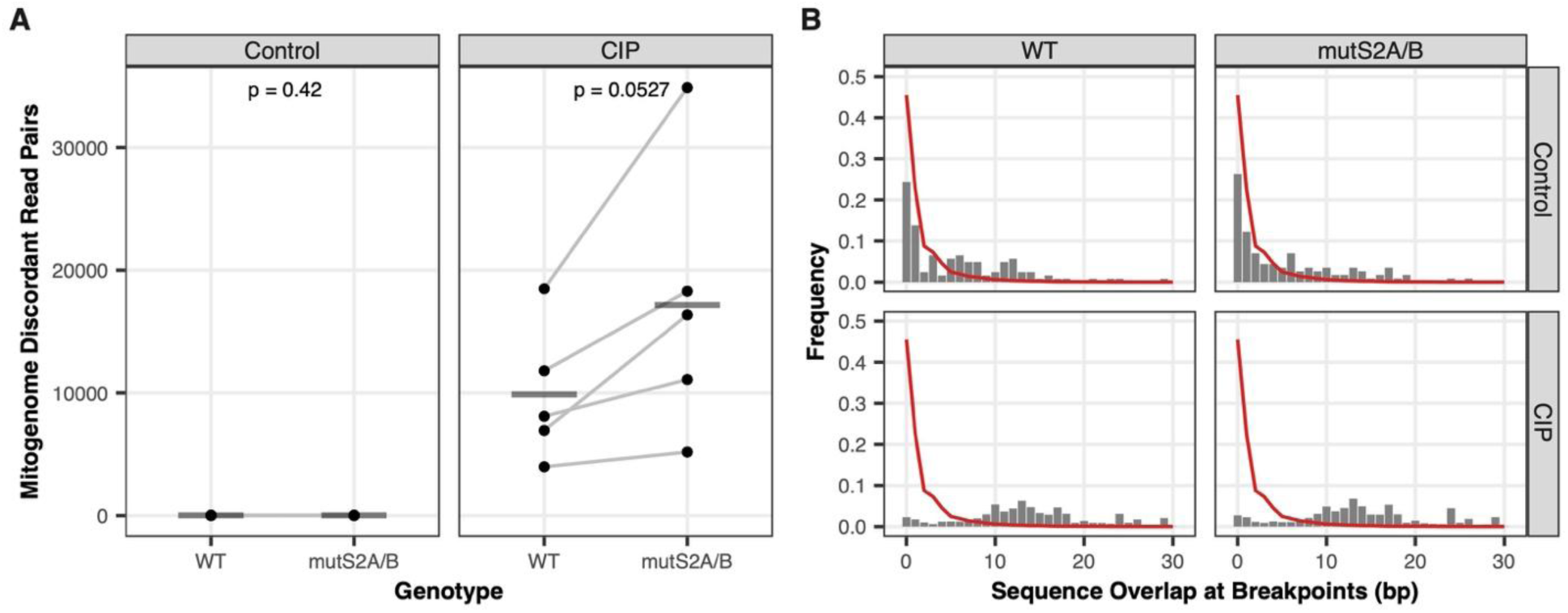
Ciprofloxacin (CIP) may disproportionately increase mitochondrial genome (mitogenome) rearrangements in *mutS2* double mutants compared to wild type. Illumina sequencing was performed on DNA from pools of seedings from individual treated with or without 0.75 μM CIP, and mitogenome rearrangements were quantified based on counts of read pairs (per million mapped) that mapped discordantly to the genome with breakpoints positioned within 1 kb of each other. (A) Points represent replicate plates containing half wild type and half mutant individuals, with lines connecting data points for samples taken from the same plate. Horizontal bars represent the mean of the five replicates. The reported p-values are based on paired *t*-test comparisons between mutant and wild type within each treatment. (B) The gray bars show the observed frequency distribution for the length of overlapping sequence similarity shared between the pairs of breakpoints for structural rearrangements in the plastid genome. The red lines show the expected distribution based on randomly sampled rearrangement positions.

**Supplemental Figure S7.**
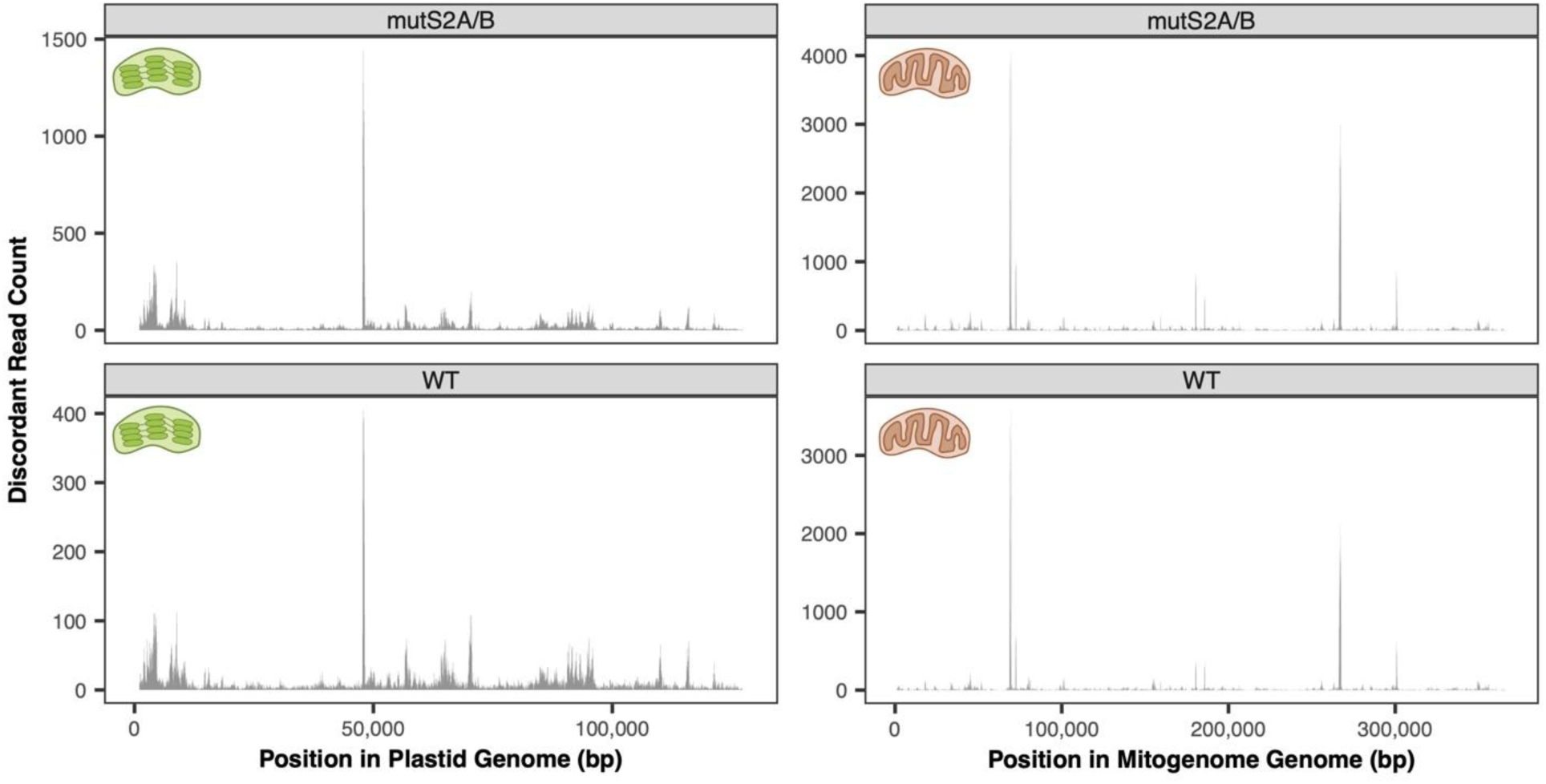
Distribution of structural rearrangement breakpoints in Arabidopsis organellar genomes resulting from ciprofloxacin treatment. Each panel shows the number of discordantly mapping Illumina sequencing reads in 50-bp sliding windows across the length of the plastid (left) and mitochondrial (right) genomes. Note that *mutS2* double mutant (top) and wild type (bottom) genotypes show nearly identical patterns of rearrangement “hotspots.” Libraries from all five replicates (see Fig. 6 and Supp. Fig. S5) were pooled for this analysis. The second copy of the plastid inverted repeat was removed for mapping purposes.

## References

1. Almagro Armenteros JJ, Salvatore M, Emanuelsson O, Winther O, von Heijne G, Elofsson A, and Nielsen H. Detecting sequence signals in targeting peptides using deep learning. Life Sci Alliance. 2019:2(5):e201900429.

2. Armarego-Marriott T, Sandoval-Ibañez O, and Kowalewska Ł. Beyond the darkness: recent lessons from etiolation and de-etiolation studies. J Exp Bot. 2020:71(4):1215–1225.

3. Barkan A and Small I. Pentatricopeptide repeat proteins in plants. Annu Rev Plant Biol. 2014:65(1):415–442.

4. Bengtson MH and Joazeiro CAP. Role of a ribosome-associated E3 ubiquitin ligase in protein quality control. Nature. 2010:467(7314):470–473.

5. Benjamini Y and Hochberg Y. Controlling the false discovery rate: A practical and powerful approach to multiple testing. J R Stat Soc Series B Stat Methodol. 1995:57(1):289–300.

6. Berdieva M, Kalinina V, Palii O, and Skarlato S. Putative MutS2 homologs in algae: More goods in shopping bag? J Mol Evol. 2024:92(6):815–833.

7. Bieri P, Leibundgut M, Saurer M, Boehringer D, and Ban N. The complete structure of the chloroplast 70S ribosome in complex with translation factor pY. EMBO J. 2017:36(4):475– 486.

8. Bulkley D, Innis CA, Blaha G, and Steitz TA. Revisiting the structures of several antibiotics bound to the bacterial ribosome. Proc Natl Acad Sci U S A. 2010:107(40):17158–17163.

9. Burby PE and Simmons LA. MutS2 Promotes Homologous Recombination in Bacillus subtilis. J Bacteriol. 2017:199(2). 10.1128/JB.00682-16

10. Callan K, Prince CR, and Feaga HA. The ribosome-associated quality control pathway supports survival in the absence of non-stop ribosome rescue factors. MBio. 2024:15(12):e0232224.

11. Cappadocia L, Maréchal A, Parent J-S, Lepage E, Sygusch J, and Brisson N. Crystal structures of DNA-Whirly complexes and their role in Arabidopsis organelle genome repair. Plant Cell. 2010:22(6):1849–1867.

12. Carrie C, Kühn K, Murcha MW, Duncan O, Small ID, O’Toole N, and Whelan J. Approaches to defining dual-targeted proteins in Arabidopsis. Plant J. 2009:57(6):1128–1139.

13. Castresana J. Selection of conserved blocks from multiple alignments for their use in phylogenetic analysis. Mol Biol Evol. 2000:17(4):540–552.

14. Cerullo F, Filbeck S, Patil PR, Hung H-C, Xu H, Vornberger J, Hofer FW, Schmitt J, Kramer G, Bukau B, et al. Bacterial ribosome collision sensing by a MutS DNA repair ATPase paralogue. Nature. 2022:603(7901):509–514.

15. Clark NL, Alani E, and Aquadro CF. Evolutionary rate covariation reveals shared functionality and coexpression of genes. Genome Res. 2012:22(4):714–720.

16. Collart MA and Weiss B. Ribosome pausing, a dangerous necessity for co-translational events. Nucleic Acids Res. 2020:48(3):1043–1055.

17. Crowe-McAuliffe C, Takada H, Murina V, Polte C, Kasvandik S, Tenson T, Ignatova Z, Atkinson GC, Wilson DN, and Hauryliuk V. Structural basis for bacterial ribosome-associated quality control by RqcH and RqcP. Mol Cell. 2021:81(1):115–126.e7.

18. Damke PP, Dhanaraju R, Marsin S, Radicella JP, and Rao DN. The nuclease activities of both the Smr domain and an additional LDLK motif are required for an efficient anti-recombination function of Helicobacter pylori MutS2. Mol Microbiol. 2015:96(6):1240–1256.

19. Diament A, Feldman A, Schochet E, Kupiec M, Arava Y, and Tuller T. The extent of ribosome queuing in budding yeast. PLoS Comput Biol. 2018:14(1):e1005951.

20. Doamekpor SK, Lee J-W, Hepowit NL, Wu C, Charenton C, Leonard M, Bengtson MH, Rajashankar KR, Sachs MS, Lima CD, et al. Structure and function of the yeast listerin (Ltn1) conserved N-terminal domain in binding to stalled 60S ribosomal subunits. Proc Natl Acad Sci U S A. 2016:113(29):E4151–60.

21. D’Orazio KN, Wu CC-C, Sinha N, Loll-Krippleber R, Brown GW, and Green R. The endonuclease Cue2 cleaves mRNAs at stalled ribosomes during No Go Decay. Elife. 2019:8:e49117.

22. Duewelhenke N, Krut O, and Eysel P. Influence on mitochondria and cytotoxicity of different antibiotics administered in high concentrations on primary human osteoblasts and cell lines. Antimicrob Agents Chemother. 2007:51(1):54–63.

23. Duval M, Dar D, Carvalho F, Rocha EPC, Sorek R, and Cossart P. HflXr, a homolog of a ribosome-splitting factor, mediates antibiotic resistance. Proc Natl Acad Sci U S A. 2018:115(52):13359–13364.

24. Eisen JA. A phylogenomic study of the MutS family of proteins. Nucleic Acids Res. 1998:26(18):4291–4300.

25. Feaga HA, Quickel MD, Hankey-Giblin PA, and Keiler KC. Human cells require non-stop ribosome rescue activity in mitochondria. PLoS Genet. 2016:12(3):e1005964.

26. Filbeck S, Cerullo F, Paternoga H, Tsaprailis G, Joazeiro CAP, and Pfeffer S. Mimicry of canonical translation elongation underlies alanine tail synthesis in RQC. Mol Cell. 2021:81(1):104–114.e6.

27. Filbeck S, Cerullo F, Pfeffer S, and Joazeiro CAP. Ribosome-associated quality-control mechanisms from bacteria to humans. Mol Cell. 2022:82(8):1451–1466.

28. Forsythe ES, Gatts TC, Lane LE, de Roux C, Berggren MJ, Rehmann EA, Zak EN, Bartel T, L’Argent LA, and Sloan DB. ERCnet: Phylogenomic prediction of interaction networks in the presence of gene duplication. Mol Biol Evol. 2025:42(5). 10.1093/molbev/msaf089

29. Forsythe ES, Grover CE, Miller ER, Conover JL, Arick MA 2nd, Chavarro MCF, Leal-Bertioli SCM, Peterson DG, Sharbrough J, Wendel JF, et al. Organellar transcripts dominate the cellular mRNA pool across plants of varying ploidy levels. Proc Natl Acad Sci U S A. 2022:119(30):e2204187119.

30. Forsythe ES, Sharbrough J, Havird JC, Warren JM, and Sloan DB. CyMIRA: The Cytonuclear Molecular Interactions Reference for Arabidopsis. Genome Biol Evol. 2019:11(8):2194– 2202.

31. Forsythe ES, Williams AM, and Sloan DB. Genome-wide signatures of plastid-nuclear coevolution point to repeated perturbations of plastid proteostasis systems across angiosperms. Plant Cell. 2021:33(4):980–997.

32. Fuchs P, Rugen N, Carrie C, Elsässer M, Finkemeier I, Giese J, Hildebrandt TM, Kühn K, Maurino VG, Ruberti C, et al. Single organelle function and organization as estimated from Arabidopsis mitochondrial proteomics. Plant J. 2020:101(2):420–441.

33. Fukui K, Inoue M, Murakawa T, Baba S, Kumasaka T, and Yano T. Structural and functional insights into the mechanism by which MutS2 recognizes a DNA junction. Structure. 2022:30(7):973–982.e4.

34. Fukui K, Kosaka H, Kuramitsu S, and Masui R. Nuclease activity of the MutS homologue MutS2 from Thermus thermophilus is confined to the Smr domain. Nucleic Acids Res. 2007:35(3):850–860.

35. Fukui K and Kuramitsu S. Structure and Function of the Small MutS-Related Domain. Mol Biol Int. 2011:2011:691735.

36. Fukui K, Masui R, and Kuramitsu S. Thermus thermophilus MutS2, a MutS paralogue, possesses an endonuclease activity promoted by MutL. J Biochem. 2004:135(3):375–384.

37. Fukui K, Murakawa T, Hino N, Kondo N, and Yano T. ATP binding controls the molecular function of bacterial MutS2 by mediating closure of the dimeric clamp structure. Structure. 2025. 10.1016/j.str.2025.03.003

38. Fukui K, Nakagawa N, Kitamura Y, Nishida Y, Masui R, and Kuramitsu S. Crystal structure of MutS2 endonuclease domain and the mechanism of homologous recombination suppression. J Biol Chem. 2008:283(48):33417–33427.

39. García-Medel PL, Baruch-Torres N, Peralta-Castro A, Trasviña-Arenas CH, Torres-Larios A, and Brieba LG. Plant organellar DNA polymerases repair double-stranded breaks by microhomology-mediated end-joining. Nucleic Acids Res. 2019:47(6):3028–3044.

40. Gawroński P, Jensen PE, Karpiński S, Leister D, and Scharff LB. Pausing of chloroplast ribosomes is induced by multiple features and is linked to the assembly of photosynthetic complexes. Plant Physiol. 2018:176(3):2557–2569.

41. Graf M, Arenz S, Huter P, Dönhöfer A, Novácek J, and Wilson DN. Cryo-EM structure of the spinach chloroplast ribosome reveals the location of plastid-specific ribosomal proteins and extensions. Nucleic Acids Res. 2017:45(5):2887–2896.

42. Greiner S, Golczyk H, Malinova I, Pellizzer T, Bock R, Börner T, and Herrmann RG. Chloroplast nucleoids are highly dynamic in ploidy, number, and structure during angiosperm leaf development. Plant J. 2020:102(4):730–746.

43. Gueneau de Novoa P and Williams KP. The tmRNA website: reductive evolution of tmRNA in plastids and other endosymbionts. Nucleic Acids Res. 2004:32(Database issue):D104–8.

44. Harrison SJ, Mott EK, Parsley K, Aspinall S, Gray JC, and Cottage A. A rapid and robust method of identifying transformed Arabidopsis thaliana seedlings following floral dip transformation. Plant Methods. 2006:2(1):19.

45. Heinemann B, Künzler P, Eubel H, Braun H-P, and Hildebrandt TM. Estimating the number of protein molecules in a plant cell: protein and amino acid homeostasis during drought. Plant Physiol. 2021:185(2):385–404.

46. Hillen HS, Lavdovskaia E, Nadler F, Hanitsch E, Linden A, Bohnsack KE, Urlaub H, and Richter-Dennerlein R. Structural basis of GTPase-mediated mitochondrial ribosome biogenesis and recycling. Nat Commun. 2021:12(1):3672.

47. Huang M, Friso G, Nishimura K, Qu X, Olinares PDB, Majeran W, Sun Q, and van Wijk KJ. Construction of plastid reference proteomes for maize and Arabidopsis and evaluation of their orthologous relationships; the concept of orthoproteomics. J Proteome Res. 2013:12(1):491–504.

48. Jacob Y, Sharkady SM, Bhardwaj K, Sanda A, and Williams KP. Function of the SmpB tail in transfer-messenger RNA translation revealed by a nucleus-encoded form. J Biol Chem. 2005:280(7):5503–5509.

49. Jarvis P and López-Juez E. Biogenesis and homeostasis of chloroplasts and other plastids. Nat Rev Mol Cell Biol. 2013:14(12):787–802.

50. Jeong E, Jo H, Kim TG, and Ban C. Characterization of multi-functional properties and conformational analysis of MutS2 from Thermotoga maritima MSB8. PLoS One. 2012:7(4):e34529.

51. Katoh K and Standley DM. MAFFT multiple sequence alignment software version 7: improvements in performance and usability. Mol Biol Evol. 2013:30(4):772–780.

52. Koller TO, Turnbull KJ, Vaitkevicius K, Crowe-McAuliffe C, Roghanian M, Bulvas O, Nakamoto JA, Kurata T, Julius C, Atkinson GC, et al. Structural basis for HflXr-mediated antibiotic resistance in Listeria monocytogenes. Nucleic Acids Res. 2022:50(19):11285–11300.

53. Langmead B and Salzberg SL. Fast gapped-read alignment with Bowtie 2. Nat Methods. 2012:9(4):357–359.

54. Lavdovskaia E, Denks K, Nadler F, Steube E, Linden A, Urlaub H, Rodnina MV, and Richter-Dennerlein R. Dual function of GTPBP6 in biogenesis and recycling of human mitochondrial ribosomes. Nucleic Acids Res. 2020:48(22):12929–12942.

55. Legen J and Schmitz-Linneweber C. Stable membrane-association of mRNAs in etiolated, greening and mature plastids. Int J Mol Sci. 2017:18(9). 10.3390/ijms18091881

56. Lin Z, Nei M, and Ma H. The origins and early evolution of DNA mismatch repair genes--multiple horizontal gene transfers and co-evolution. Nucleic Acids Res. 2007:35(22):7591–7603.

57. Lytvynenko I, Paternoga H, Thrun A, Balke A, Müller TA, Chiang CH, Nagler K, Tsaprailis G, Anders S, Bischofs I, et al. Alanine tails signal proteolysis in bacterial ribosome-associated quality control. Cell. 2019:178(1):76–90.e22.

58. Majeran W, Friso G, Asakura Y, Qu X, Huang M, Ponnala L, Watkins KP, Barkan A, and van Wijk KJ. Nucleoid-enriched proteomes in developing plastids and chloroplasts from maize leaves: a new conceptual framework for nucleoid functions. Plant Physiol. 2012:158(1):156–189.

59. Malik HS and Henikoff S. Dual recognition-incision enzymes might be involved in mismatch repair and meiosis. Trends Biochem Sci. 2000:25(9):414–418.

60. Maréchal A and Brisson N. Recombination and the maintenance of plant organelle genome stability. New Phytol. 2010:186(2):299–317.

61. Martin M. Cutadapt removes adapter sequences from high-throughput sequencing reads. EMBnet J. 2011:17(1):10.

62. Mehrez M, Lecampion C, Ke H, Gorsane F, and Field B. Insights into the function of the chloroplastic ribosome-associated GTPase high frequency of lysogenization X in Arabidopsis thaliana. Plant Direct. 2024:8(1):e559.

63. Meinke DW. Genome-wide identification of EMBRYO-DEFECTIVE (EMB) genes required for growth and development in Arabidopsis. New Phytol. 2020:226(2):306–325.

64. Mi H, Muruganujan A, Huang X, Ebert D, Mills C, Guo X, and Thomas PD. Protocol Update for large-scale genome and gene function analysis with the PANTHER classification system (v.14.0). Nat Protoc. 2019:14(3):703–721.

65. Miller-Messmer M, Kühn K, Bichara M, Le Ret M, Imbault P, and Gualberto JM. RecA-dependent DNA repair results in increased heteroplasmy of the Arabidopsis mitochondrial genome. Plant Physiol. 2012:159(1):211–226.

66. Moore SD and Sauer RT. Ribosome rescue: tmRNA tagging activity and capacity in Escherichia coli: tmRNA tagging inE. coli. Mol Microbiol. 2005:58(2):456–466.

67. Moore SD and Sauer RT. The tmRNA system for translational surveillance and ribosome rescue. Annu Rev Biochem. 2007:76(1):101–124.

68. Mulo P, Pursiheimo S, Hou C-X, Tyystjärvi T, and Aro E-M. Multiple effects of antibiotics on chloroplast and nuclear gene expression. Funct Plant Biol. 2003:30(11):1097–1103.

69. Nadler F, Lavdovskaia E, and Richter-Dennerlein R. Maintaining mitochondrial ribosome function: The role of ribosome rescue and recycling factors. RNA Biol. 2022:19(1):117–131.

70. Nagao M, Tsuchiya F, Motohashi R, and Abo T. Ribosome rescue activity of an Arabidopsis thaliana ArfB homolog. Genes Genet Syst. 2020:95(3):119–131.

71. Nanjaraj Urs AN, Lasehinde V, Kim L, McDonald E, Yan LL, and Zaher HS. Inability to rescue stalled ribosomes results in overactivation of the integrated stress response. J Biol Chem. 2024:300(5):107290.

72. Nishimura K, Kato Y, and Sakamoto W. Essentials of proteolytic machineries in chloroplasts. Mol Plant. 2017:10(1):4–19.

73. Obayashi T, Hibara H, Kagaya Y, Aoki Y, and Kinoshita K. ATTED-II v11: A plant gene coexpression database using a sample balancing technique by subagging of principal components. Plant Cell Physiol. 2022:63(6):869–881.

74. Oelmüller R, Levitan I, Bergfeld R, Rajasekhar VK, and Mohr H. Expression of nuclear genes as affected by treatments acting on the plastids. Planta. 1986:168(4):482–492.

75. Ogata H, Ray J, Toyoda K, Sandaa R-A, Nagasaki K, Bratbak G, and Claverie J-M. Two new subfamilies of DNA mismatch repair proteins (MutS) specifically abundant in the marine environment. ISME J. 2011:5(7):1143–1151.

76. Olinares PDB, Ponnala L, and van Wijk KJ. Megadalton complexes in the chloroplast stroma of Arabidopsis thaliana characterized by size exclusion chromatography, mass spectrometry, and hierarchical clustering. Mol Cell Proteomics. 2010:9(7):1594–1615.

77. Parent J-S, Lepage E, and Brisson N. Divergent roles for the two PolI-like organelle DNA polymerases of Arabidopsis. Plant Physiol. 2011:156(1):254–262.

78. Park EN, Mackens-Kiani T, Berhane R, Esser H, Erdenebat C, Burroughs AM, Berninghausen O, Aravind L, Beckmann R, Green R, et al. B. subtilis MutS2 splits stalled ribosomes into subunits without mRNA cleavage. EMBO J. 2024:43(4):484–506.

79. Pinto AV, Mathieu A, Marsin S, Veaute X, Ielpi L, Labigne A, and Radicella JP. Suppression of homologous and homeologous recombination by the bacterial MutS2 protein. Mol Cell. 2005:17(1):113–120.

80. Pipitone R, Eicke S, Pfister B, Glauser G, Falconet D, Uwizeye C, Pralon T, Zeeman SC, Kessler F, and Demarsy E. A multifaceted analysis reveals two distinct phases of chloroplast biogenesis during de-etiolation in Arabidopsis. Elife. 2021:10. 10.7554/eLife.62709

81. Puthiyaveetil S, McKenzie SD, Kayanja GE, and Ibrahim IM. Transcription initiation as a control point in plastid gene expression. Biochim Biophys Acta Gene Regul Mech. 2021:1864(3):194689.

82. Rhee SY, Beavis W, Berardini TZ, Chen G, Dixon D, Doyle A, Garcia-Hernandez M, Huala E, Lander G, Montoya M, et al. The Arabidopsis Information Resource (TAIR): a model organism database providing a centralized, curated gateway to Arabidopsis biology, research materials and community. Nucleic Acids Res. 2003:31(1):224–228.

83. Richter R, Rorbach J, Pajak A, Smith PM, Wessels HJ, Huynen MA, Smeitink JA, Lightowlers RN, and Chrzanowska-Lightowlers ZM. A functional peptidyl-tRNA hydrolase, ICT1, has been recruited into the human mitochondrial ribosome. EMBO J. 2010:29(6):1116–1125.

84. Rocha DC, da Silva Rocha C, Tavares DS, de Morais Calado SL, and Gomes MP. Veterinary antibiotics and plant physiology: An overview. Sci Total Environ. 2021:767(144902):144902.

85. Rudra P, Hurst-Hess KR, Cotten KL, Partida-Miranda A, and Ghosh P. Mycobacterial HflX is a ribosome splitting factor that mediates antibiotic resistance. Proc Natl Acad Sci U S A. 2020:117(1):629–634.

86. Sachadyn P. Conservation and diversity of MutS proteins. Mutat Res. 2010:694(1–2):20–30.

87. Saito K, Kratzat H, Campbell A, Buschauer R, Burroughs AM, Berninghausen O, Aravind L, Green R, Beckmann R, and Buskirk AR. Ribosome collisions induce mRNA cleavage and ribosome rescue in bacteria. Nature. 2022:603(7901):503–508.

88. Sakamoto W. Protein degradation machineries in plastids. Annu Rev Plant Biol. 2006:57(1):599– 621.

89. Schatz-Daas D, Fertet A, Lotfi F, and Gualberto JM. Assessment of mitochondrial DNA copy number, stability, and repair in Arabidopsis. Methods Mol Biol. 2022:2363:301–319.

90. Shen PS, Park J, Qin Y, Li X, Parsawar K, Larson MH, Cox J, Cheng Y, Lambowitz AM, Weissman JS, et al. Protein synthesis. Rqc2p and 60S ribosomal subunits mediate mRNA-independent elongation of nascent chains. Science. 2015:347(6217):75–78.

91. Sloan DB, Broz AK, Kuster SA, Muthye V, Peñafiel-Ayala A, Marron JR, Lavrov DV, and Brieba LG. Expansion of the MutS gene family in plants. 2024:koae277.

92. Small I, Peeters N, Legeai F, and Lurin C. Predotar: A tool for rapidly screening proteomes for N-terminal targeting sequences. Proteomics. 2004:4(6):1581–1590.

93. Snowden T, Acharya S, Butz C, Berardini M, and Fishel R. hMSH4-hMSH5 recognizes Holliday Junctions and forms a meiosis-specific sliding clamp that embraces homologous chromosomes. Mol Cell. 2004:15(3):437–451.

94. Sperschneider J, Catanzariti A-M, DeBoer K, Petre B, Gardiner DM, Singh KB, Dodds PN, and Taylor JM. LOCALIZER: subcellular localization prediction of both plant and effector proteins in the plant cell. Sci Rep. 2017:7(1):44598.

95. Stamatakis A. RAxML version 8: a tool for phylogenetic analysis and post-analysis of large phylogenies. Bioinformatics. 2014:30(9):1312–1313.

96. Suárez-Rivero JM, Pastor-Maldonado CJ, Povea-Cabello S, Álvarez-Córdoba M, Villalón-García I, Talaverón-Rey M, Suárez-Carrillo A, Munuera-Cabeza M, and Sánchez-Alcázar JA. Mitochondria and antibiotics: For good or for evil? Biomolecules. 2021:11(7):1050.

97. Takada H, Crowe-McAuliffe C, Polte C, Sidorova ZY, Murina V, Atkinson GC, Konevega AL, Ignatova Z, Wilson DN, and Hauryliuk V. RqcH and RqcP catalyze processive poly-alanine synthesis in a reconstituted ribosome-associated quality control system. Nucleic Acids Res. 2021:49(14):8355–8369.

98. Thomas PD, Ebert D, Muruganujan A, Mushayahama T, Albou L-P, and Mi H. PANTHER: Making genome-scale phylogenetics accessible to all. Protein Sci. 2022:31(1):8–22.

99. Waese J, Fan J, Pasha A, Yu H, Fucile G, Shi R, Cumming M, Kelley LA, Sternberg MJ, Krishnakumar V, et al. EPlant: Visualizing and exploring multiple levels of data for hypothesis generation in plant biology. Plant Cell. 2017:29(8):1806–1821.

100. Wang G, Alamuri P, Humayun MZ, Taylor DE, and Maier RJ. The Helicobacter pylori MutS protein confers protection from oxidative DNA damage. Mol Microbiol. 2005:58(1):166–176.

101. Wang G and Maier RJ. Molecular basis for the functions of a bacterial MutS2 in DNA repair and recombination. DNA Repair. 2017:57:161–170.

102. Wang X, Ryu D, Houtkooper RH, and Auwerx J. Antibiotic use and abuse: a threat to mitochondria and chloroplasts with impact on research, health, and environment. Bioessays. 2015:37(10):1045–1053.

103. Westrich LD, Gotsmann VL, Herkt C, Ries F, Kazek T, Trösch R, Armbruster L, Mühlenbeck JS, Ramundo S, Nickelsen J, et al. The versatile interactome of chloroplast ribosomes revealed by affinity purification mass spectrometry. Nucleic Acids Res. 2021:49(1):400–415.

104. van Wijk KJ. Protein maturation and proteolysis in plant plastids, mitochondria, and peroxisomes. Annu Rev Plant Biol. 2015:66(1):75–111.

105. van Wijk KJ. Intra-chloroplast proteases: A holistic network view of chloroplast proteolysis. Plant Cell. 2024:36(9):3116–3130.

106. van Wijk KJ, Leppert T, Sun Q, Boguraev SS, Sun Z, Mendoza L, and Deutsch EW. The Arabidopsis PeptideAtlas: Harnessing worldwide proteomics data to create a comprehensive community proteomics resource. Plant Cell. 2021:33(11):3421–3453.

107. van Wijk KJ, Leppert T, Sun Z, and Deutsch EW. Does the ubiquitination degradation pathway really reach inside of the chloroplast? A re-evaluation of mass spectrometry-based assignments of ubiquitination. J Proteome Res. 2023:22(6):2079–2091.

108. Williams KP and Bartel DP. The tmRNA website. Nucleic Acids Res. 1998.

109. Wilson DN. Ribosome-targeting antibiotics and mechanisms of bacterial resistance. Nat Rev Microbiol. 2014:12(1):35–48.

110. Xiao Y, Xiong T, Meng X, Yu D, Xiao Z, and Song L. Different influences on mitochondrial function, oxidative stress and cytotoxicity of antibiotics on primary human neuron and cell lines: XIAO et al. J Biochem Mol Toxicol. 2019:33(4):e22277.

111. Xiong H-B, Pan H-M, Long Q-Y, Wang Z-Y, Qu W-T, Mei T, Zhang N, Xu X-F, Yang Z-N, and Yu Q-B. AtNusG, a chloroplast nucleoid protein of bacterial origin linking chloroplast transcriptional and translational machineries, is required for proper chloroplast gene expression in Arabidopsis thaliana. Nucleic Acids Res. 2022:50(12):6715–6734.

112. Yan LL and Zaher HS. Ribosome quality control antagonizes the activation of the integrated stress response on colliding ribosomes. Mol Cell. 2021:81(3):614–628.e4.

113. Yang M, Derbyshire MK, Yamashita RA, and Marchler-Bauer A. NCBI’s Conserved Domain Database and tools for protein domain analysis. Curr Protoc Bioinformatics. 2020:69(1):e90.

114. Yoon Y-E, Cho HM, Bae D-W, Lee SJ, Choe H, Kim MC, Cheong MS, and Lee YB. Erythromycin treatment of Brassica campestris seedlings impacts the photosynthetic and protein synthesis pathways. Life (Basel). 2020:10(12):311.

115. Yu G. Using ggtree to Visualize Data on Tree-Like Structures. Curr Protoc Bioinformatics. 2020:69(1):e96.

116. Zampini É, Lepage É, Tremblay-Belzile S, Truche S, and Brisson N. Organelle DNA rearrangement mapping reveals U-turn-like inversions as a major source of genomic instability in Arabidopsis and humans. Genome Res. 2015:25(5):645–654.

117. Zhang H, Xu Q, Lu M, Xu X, Wang Y, Wang L, Zhao Y, and Hua Y. Structural and functional studies of MutS2 from Deinococcus radiodurans. DNA Repair. 2014:21:111–119.

118. Zhang J, Ruhlman TA, Sabir JSM, Blazier JC, Weng M-L, Park S, and Jansen RK. Coevolution between nuclear-encoded DNA replication, recombination, and repair genes and Plastid genome complexity. Genome Biol Evol. 2016:8(3):622–634.

119. Zhang X, Henriques R, Lin S-S, Niu Q-W, and Chua N-H. Agrobacterium-mediated transformation of Arabidopsis thaliana using the floral dip method. Nat Protoc. 2006:1(2):641–646.

120. Zhang Y, Mandava CS, Cao W, Li X, Zhang D, Li N, Zhang Y, Zhang X, Qin Y, Mi K, et al. HflX is a ribosome-splitting factor rescuing stalled ribosomes under stress conditions. Nat Struct Mol Biol. 2015:22(11):906–913.

121. Zhang Y, Tian L, and Lu C. Chloroplast gene expression: Recent advances and perspectives. Plant Commun. 2023:4(5):100611.

122. Zhao T, Chen Y-M, Li Y, Wang J, Chen S, Gao N, and Qian W. Disome-seq reveals widespread ribosome collisions that promote cotranslational protein folding. Genome Biol. 2021:22(1):16.

123. Zhou W, Lu Q, Li Q, Wang L, Ding S, Zhang A, Wen X, Zhang L, and Lu C. PPR-SMR protein SOT1 has RNA endonuclease activity. Proc Natl Acad Sci U S A. 2017:114(8):E1554–E1563.

124. Zoschke R and Bock R. Chloroplast translation: Structural and functional organization, operational control, and regulation. Plant Cell. 2018:30(4):745–770.

